# A localization screen reveals translation factories and widespread co-translational RNA targeting

**DOI:** 10.1101/2020.05.20.106989

**Authors:** Racha Chouaib, Adham Safieddine, Xavier Pichon, Arthur Imbert, Oh Sung Kwon, Aubin Samacoits, Abdel-Meneem Traboulsi, Marie-Cécile Robert, Nikolay Tsanov, Emeline Coleno, Ina Poser, Christophe Zimmer, Anthony Hyman, Hervé Le Hir, Kazem Zibara, Marion Peter, Florian Mueller, Thomas Walter, Edouard Bertrand

## Abstract

Local translation allows a spatial control of gene expression. Here, we performed a dual protein/mRNA localization screen, using smFISH on 523 human cell lines expressing GFP-tagged genes. A total of 32 mRNAs displayed specific cytoplasmic localizations, and we observed local translation at unexpected locations, including cytoplasmic protrusions, cell edges, endosomes, Golgi, the nuclear envelope and centrosomes, the latter being cell cycle dependent. Quantitation of mRNA distribution and automatic pattern classification revealed a high degree of localization heterogeneity between cells. Surprisingly, mRNA localization frequently required ongoing translation, indicating widespread co-translational RNA targeting. Interestingly, while P-body accumulation was frequent (15 mRNAs), four mRNAs accumulated in foci that were distinct structures. These foci lacked the mature protein, but nascent polypeptide imaging showed that they were specialized translation factories. For β-catenin, foci formation was regulated by Wnt, relied on APC-dependent polysome aggregation, and led to nascent protein degradation. Thus, translation factories uniquely regulate nascent protein metabolism and create a fine granular compartmentalization of translation.

## Introduction

Most mRNAs are distributed randomly throughout the cytoplasm, but some localize to specific subcellular areas (Blower, 2013; Bovaird et al., 2018; Eliscovich and Singer, 2017; Jung et al., 2014 for reviews). This phenomenon is linked to either RNA metabolism, when untranslated mRNAs are stored in P-bodies or other cellular structures (Hubstenberger et al., 2017); or to protein metabolism, when a protein is synthesized locally. Local translation has been observed from bacteria and yeast to humans (Blower, 2013; Jung et al., 2014; Eliscovich and Singer, 2017; Bovaird et al., 2018). It is commonly involved in the delivery of mature proteins to specific cellular compartments, and this is involved in many processes. For instance, it contributes to patterning and cell fate determination during metazoan development, mainly through asymmetric cell division (Melton, 1987; Driever and Nüsslein-Volhard, 1988). In Xenopus embryos, local translation of cyclin B mRNAs at the mitotic spindle is also believed to be important for the rapid cell division cycles occurring during early embryogenesis (Groisman et al., 2000). In mammals, mRNA localization is involved in cell polarization and motility, mainly through the localization of actin and related mRNAs at the leading edge (Lawrence and Singer, 1986), and it is also involved in axonal growth and synaptic plasticity of neurons (Van Driesche and Martin, 2018). Importantly, local translation can also be linked to the metabolism of the nascent peptide rather than to directly localize the mature protein. For instance, translation of secreted proteins at the endoplasmic reticulum (ER) allows nascent proteins to translocate through the membrane to reach the ER lumen (Aviram and Schuldiner, 2017). Translation of mRNAs at specific sites may also be important for the assembly of protein complexes (Pichon et al., 2016), or to avoid the deleterious effects of releasing free proteins at inappropriate places (Müller et al., 2013).

RNA localization can be accomplished through several mechanisms. In the case of secreted proteins, the nascent peptide serves as a targeting signal, via the signal recognition particle (SRP) and its receptor on the ER (Aviram and Schuldiner, 2017). In most other cases, targeting is an RNA-driven process (Blower, 2013; Jung et al., 2014; Eliscovich and Singer, 2017; Bovaird et al., 2018 for reviews). Localized mRNAs often contain a zip-code sequence, frequently located whitin their 3’-UTR, which is necessary and sufficient to transport them to their destination. The zipcode is recognized by one or several RNA-binding proteins (RBP), and it drives the formation of a transport complex sometimes called locasome. This complex can be transported by centrosomes, endosomal vesicles or other cellular structures (Blower, 2013; Jung et al., 2014; Eliscovich and Singer, 2017; Bovaird et al., 2018). However, direct transport on the cytoskeleton by molecular motors is a frequent mechanism (Blower, 2013; Bovaird et al., 2018; Eliscovich and Singer, 2017; Jung et al., 2014). Once at destination, an anchoring mechanism may limit diffusion away from the target site. Alternatives to these transport mechanisms include diffusion and trapping at specific locations, and degradation coupled to local RNA stabilization.

Localized mRNAs are often subjected to a spatial control of translation (Besse and Ephrussi, 2008). In the case of Ash1 mRNA in yeast and β-actin mRNA in neurons, translation is repressed during transport and is activated at their final location by phosphorylation-dependent mechanisms (Hüttelmaier et al., 2005; Paquin et al., 2007). This spatial regulation of translation provides an additional layer of control ensuring that mRNAs are translated only at the desired location.

The first locally-translated mRNAs were found by chance or using a candidate approach. Purification of cellular structures and localized RBPs have significantly increased the number of known localized mRNAs (Blower, 2013; Jung et al., 2014; Eliscovich and Singer, 2017; Bovaird et al., 2018). However, only specific compartments or RBPs were examined and we currently lack a global view of local translation in the entire cellular space or at the genomic level. Few reports described attempts to characterize mRNA localization in a systematic manner. A pioneering study in Drosophila used whole-mount fluorescent *in situ* hybridization to analyze the localization of more than 2000 mRNAs (Lécuyer et al., 2007). As many as 71% of them had a non-random distribution, and a range of new localization patterns were observed. More recent reports confirmed that RNA localization is widespread during Drosophila development (Jambor et al., 2015; Wilk et al., 2016). However, it is not known whether this is also true in other organisms and particularly in humans. Few recent studies addressed this question in cell lines using the more sensitive single molecule FISH technique (smFISH). Several thousands of mRNAs were analyzed, which showed a correlation of intracellular mRNA distribution with gene annotation (Battich et al., 2013; Chen et al., 2015; Eng et al., 2019; Xia et al., 2019). Specifically, these studies identified three groups of localized mRNAs, in the perinuclear area, the mitochondria and the cell periphery, the latter being possibly linked to actin metabolism (Chen et al., 2015).

These studies provided information on RNA localization but did not directly investigate local translation as the encoded proteins were not detected. Thus, we still lack a good understanding of the various functions played by local translation at the cellular level. In this study, we developed a smFISH screen to specifically address this issue. Using a set of 523 GFP-tagged cell lines spanning a variety of cellular functions, and an approach that allows simultaneous visualization of mRNA and proteins, we found that local translation occurs at various unanticipated locations. In particular, we discovered specialized translation factories, where specific mRNAs are translated. These factories are remarkable in that they provide a unique mean to regulate the metabolism of nascent proteins and also create a fine granular compartmentalization of translation.

## Results

### BAC TransgeneOmics allow dual protein/RNA localization screens

In order to simultaneously visualize mRNAs with their encoded proteins, we based our screen on a library of HeLa cell lines, each containing a bacterial artificial chromosome (BAC) stably integrated in their genome (Poser et al., 2008). Each BAC contains a GFP-tagged gene harbouring all its regulatory sequences (promoter, enhancers, introns, 5’ and 3’ UTRs; Figure 1A). The resulting mRNAs are thus identical to the endogenous molecules in terms of sequence and isoform diversity, except for the added tag. Previous studies showed that such tagged genes are expressed at near endogenous levels and with the proper spatio-temporal pattern (Poser et al., 2008). Since the tagged mRNAs contain all the regulatory sequences, we hypothesized that they would localize like the endogenous ones, provided that the tag does not interfere with localization. Using BACs offers two advantages. First, a single smFISH probe set against the GFP sequence is sufficient to detect all the studied mRNAs. Second, using mild hybridization conditions, GFP fluorescence can be detected together with the smFISH signal (Fusco et al., 2003), and thus both the mRNA and the encoded protein can be detected in the same cell. Genes in the BAC collection are tagged at either ends. N-terminally tagged proteins (N-FLAP in Table S1 and S2) have only the GFP tag since the selectable marker is placed within an intron, whereas C-terminally tagged proteins (C-LAP in Table S1 and S2) carry a GFP-IRES-Neo tag that allows both visualization and selection (Poser et al., 2008). We developed a set of 19 fluorescent oligonucleotide probes against the GFP tag and an additional 25 probes against the IRES-neo sequence (Figure 1A, Table S3). Following smFISH with these probes, no signal was detected in the parental HeLa cells while isolated spots corresponding to single mRNAs were visible in the TERF1-GFP or GFP-VAPA BAC cells (Figure 1B). The signal was more intense for the C-terminally tagged mRNAs because of the higher number of probes. Importantly, the GFP signal was visible after smFISH, demonstrating the feasibility of the approach.

**Figure 1.**
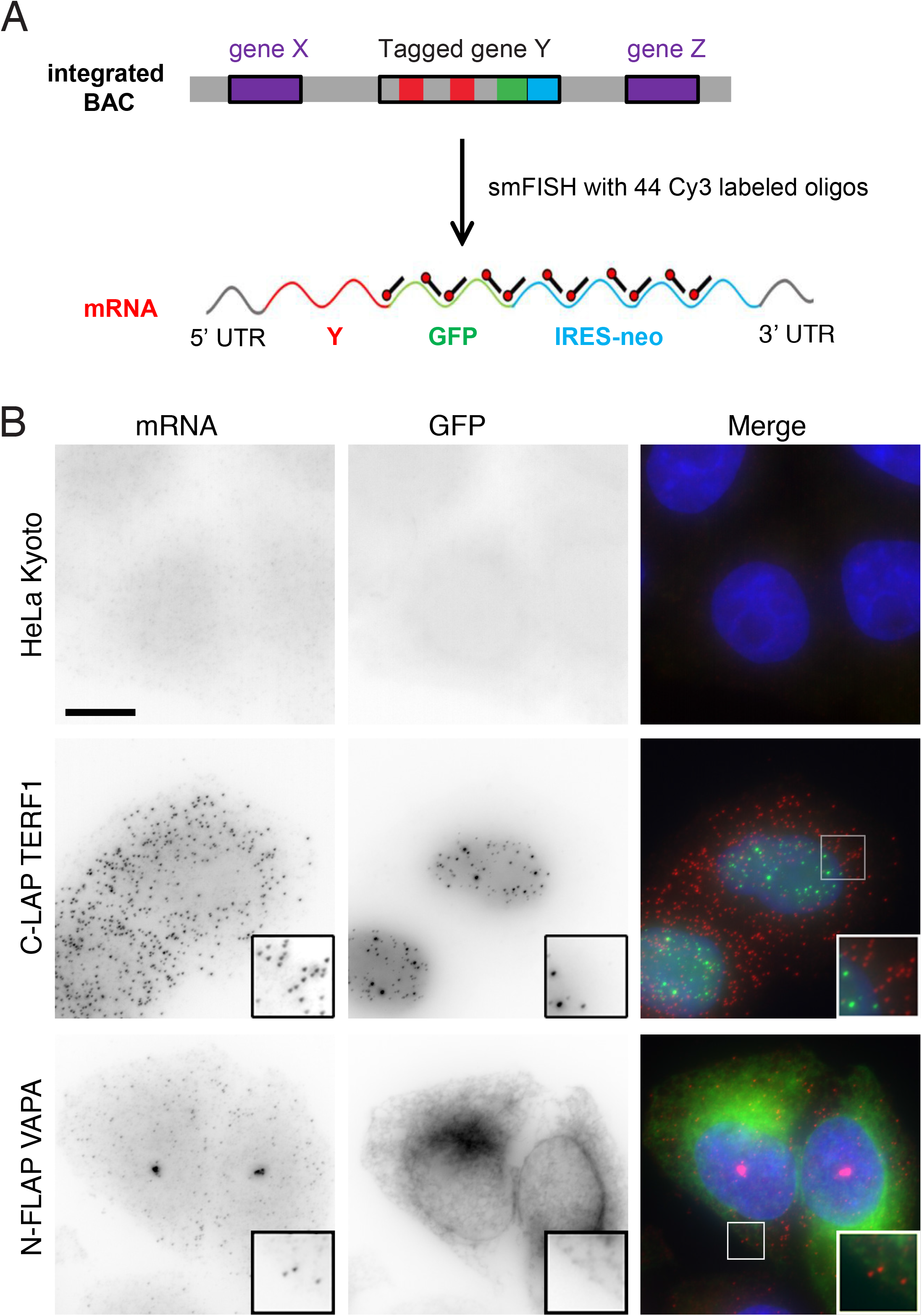
A dual protein/mRNA localization screen using GFP-tagged BACs. A-Schematic of the BAC screen by smFISH. The schematic depicts the integrated BAC with the tagged gene (top; exons in red), which transcribes a GFP-tagged mRNA with all its regulatory sequences (bottom; 5’ and 3’ UTR). The location of the probes is indicated (black pins with red head), and X and Z represent neighboring genes. B-The BACs allow simultaneous detection of mRNAs and their encoded proteins. Panels are micrographs of HeLa cells expressing the indicated BAC, or no BAC as control (top), and hybridized with Cy3-labelled probes against the GFP-IRES-Neo sequence. Red and left: signal from the smFISH probes; green and middle: signal from the GFP channel; blue: DAPI staining. Scale bar: 10 microns. Insets: zoom of the boxed areas.

### Screening transcripts encoding motors reveals six localized mRNAs and three patterns

Molecular motors transport cargos to different cellular destinations and accumulate themselves at various cellular locations. The BAC collection contains all the human genes coding for kinesins and myosins (59 and 44, respectively; Maliga et al., 2013), as well as 7 out of 11 dynein subunits. These gene families were screened to test whether some mRNAs would localize to specific cellular areas. We hypothesized that this could reveal a local function of motor proteins, for instance as a mean to transport cargos from a particular location. Most motor mRNAs localized randomly throughout the cytoplasm (Table S1). However, 6 displayed atypical localizations, with three distinct patterns (Figure 2 and Table S2). In the first, referred to as “*protrusion*”, mRNAs accumulated in cytoplasmic extensions. This was observed for three kinesins (KIF1C, KIF4A and KIF5B), the dynein subunit DYNLL2 and the myosin MYH3 (Figure 2A). Second, MYH3 mRNAs additionally accumulated inside the nucleus, a pattern referred to as “*intranuclear*”. The third pattern, referred to as “*fOci*”, was observed for the dynein heavy chain mRNA (DYNC1H1). This mRNA localized throughout the cytoplasm but aggregated in some bright structures containing several mRNA molecules (Figure 2B), as reported recently (Pichon et al., 2016). Surprisingly, the proteins encoded by some of these localized mRNAs did not appear strongly enriched at the site of mRNA accumulation. GFP-KIF4A was mostly nuclear with only a faint staining at the cell periphery, while GFP-tagged KIF5B, DYNLL2 and DYNC1H1 proteins localized throughout the cytoplasm without a specific enrichment in protrusions or foci (Figure 2). Co-localization was only observed for KIF1C-GFP, where both the mRNA and the GFP-tagged protein accumulated in cytoplasmic protrusions. This suggested that the KIF1C-GFP mRNAs were translated locally.

**Figure 2.**
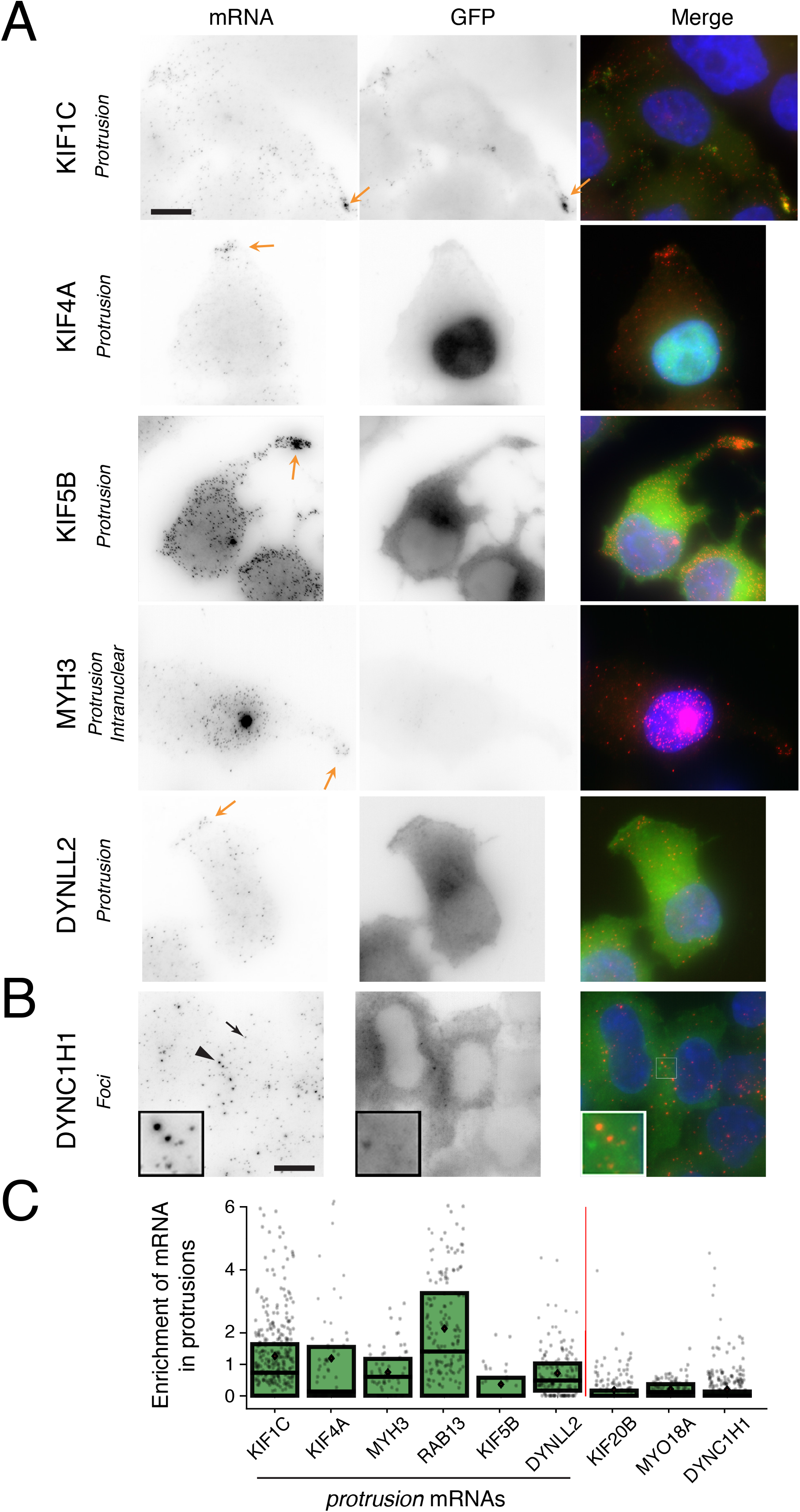
Localization of the mRNAs encoding molecular motors. A-Transcripts localizing to cytoplasmic protrusions. Legends as in Figure 1B. The name of the BAC tagged gene is on the left. Arrows indicate the mRNA accumulation at cell extensions. Scale bar: 10 microns. B-mRNAs accumulating in cytoplasmic foci. Arrow: a single mRNA; arrowhead: a foci containing multiple mRNAs. Inset: zoom of the boxed area. Scale bar: 10 microns. C-Quantification of the amount mRNA accumulating in cytoplasmic protrusions. The boxplot depicts the enrichment of mRNAs in cytoplasmic protrusion, obtained by dividing the fraction of mRNA in protrusion by the fraction of the protrusion surface. Each cell is a spot; the box corresponds to the second and third quartiles; the black bar is the median and the red diamond the mean.

To confirm the BAC results, endogenous mRNAs were analyzed using smiFISH, an inexpensive variant of smFISH (Tsanov et al., 2016). KIF1C mRNAs accumulated in cytoplasmic protrusions in all the examined cell lines, including human HeLa cells as well as mouse 3T3 fibroblasts and C2C12 myoblasts (Figure S1A). Furthermore, in differentiated SH-SY5Y neuronal cells, KIF1C mRNA localized in dendritic-like outgrowths, away from the cell body, while a control CRM1 mRNA remained in the soma (Figure S1B). Likewise, DYNC1H1 mRNAs formed foci in all examined cell lines (Figure S1A). MYH3 mRNA was not expressed in HeLa, 293HEK and 3T3, but localized in cellular protrusions in C2C12 cells, as well as in dendritic-like processes of SH-SY5Y neuronal cells. In contrast, KIF5B mRNA localization varied with the cell line. This mRNA was present in neuronal dendrite-like processes of SH-SY5Y cells, but localized only weakly in HeLa or C2C12 lines (Figure S1, data not shown).

To provide a quantitative view of mRNA localization, we developed tools to identify the cytoplasmic protrusions and we measured both the fraction of cytoplasmic mRNAs located in this compartment and their enrichment as compared to a uniform distribution of molecules (see STAR Methods). This revealed a high heterogeneity of mRNA localization in protrusions, with values varying from 0 to 60% of the molecules depending on the cell (Figure S2A). On average, the enrichment in protrusions of KIF1C, KIF4A, DYNLL2 and MYH3 mRNAs was several fold higher than for control motor mRNAs annotated as random (KIF20B, MYO18A; Figure 2C). It was however lower than for RAB13 mRNA that we used as a positive control (Mili et al., 2008). Next, we determined whether KIF1C, KIF5B, MYH3 and KIF4A mRNAs accumulated in the same protrusions. Two-color smFISH against the endogenous KIF1C mRNAs and the B AC-tagged mRNAs revealed that this was indeed the case (Figure S2B). Taken together, these results validate our BAC approach for simultaneously screening mRNA and protein localization. We identified six localized mRNAs and three localization patterns, with KIF1C likely being locally translated in cytoplasmic protrusions.

### Systematic screening of BAC-tagged mRNAs reveals a total of six cytoplasmic localization patterns

We then analyzed 411 BAC-tagged genes involved in a variety of biological processes (Table S1). The results revealed that 26 mRNAs displayed a specific localization in the cytoplasm (about 6%). Note that some localized mRNAs may have been missed if the tag interfered with localization, or if the pattern was too subtle to be annotated manually. The localized mRNAs were classified into 6 patterns (Figure 3 and S3 for larger fields of view; Table 1 and S2). In addition to the previous patterns (‘*protrusion*” and “*foci*”), some mRNAs accumulated at the nuclear envelope (“*nuclear edge*”), around the cell periphery (“*cell edge*”), or had a tendency to localize towards the nucleus, often in a polarized manner (“*perinuclear*”; Figure 3B and S3, Table S2). Interestingly, we also found three mRNAs that accumulated at centrosomes during cell division (“*mitotic*”; HMMR, ASPM and NUMA1; Figure 4). Overall, accumulation of mRNAs in foci was the most frequent pattern (19 out of the total of 32 localized mRNAs; Table 1 and S2). To determine whether any of these foci were P-bodies, we imaged mRNAs together with a P-body marker. Indeed, foci of 15 mRNAs colocalized with P-bodies (Figure S4), while the remaining four were distinct structures (BUB1, DYNC1H1, CTNNB1/β-catenin and ASPM mRNA; Figure S5A).

**Figure 3.**
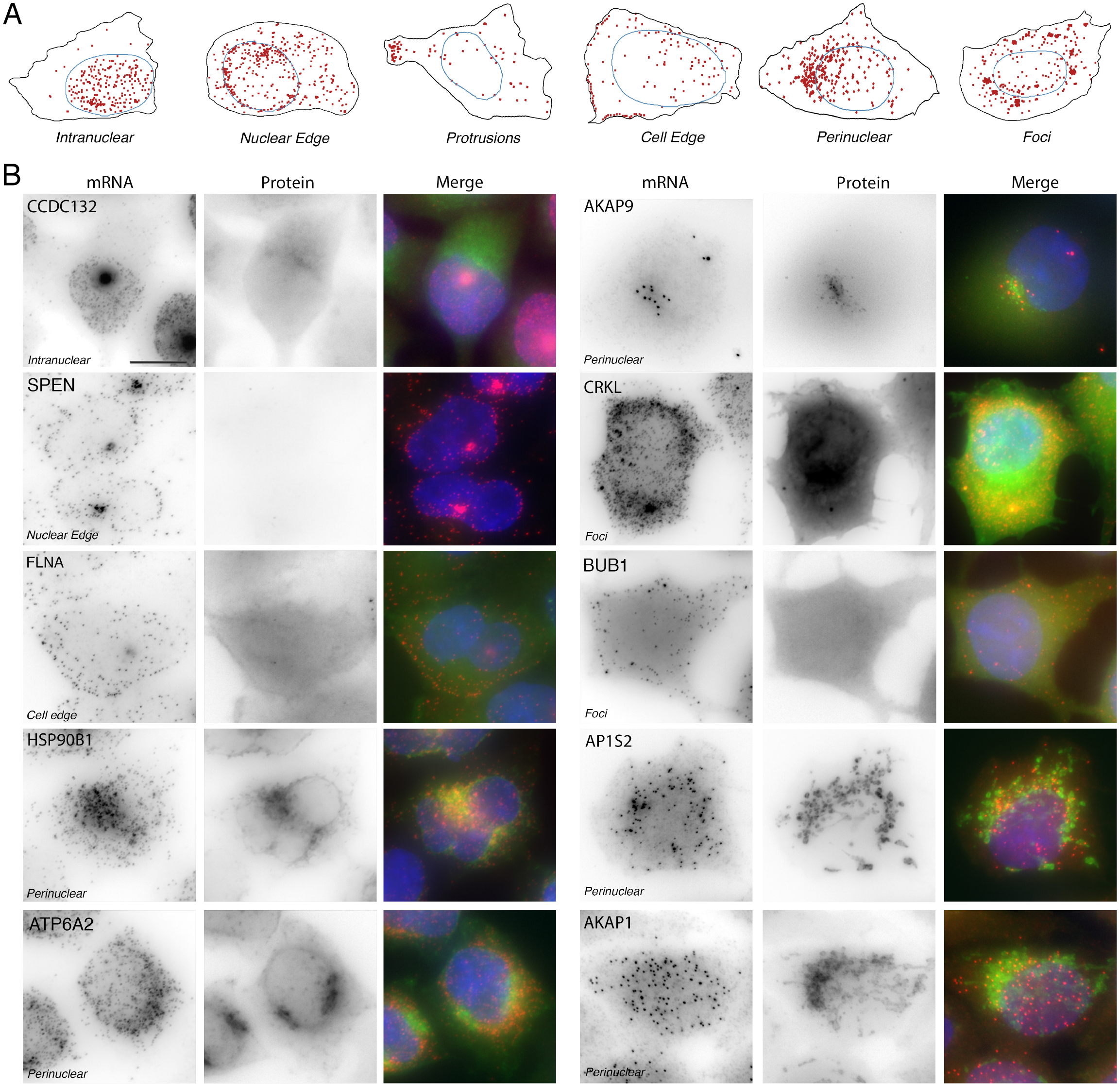
Examples of localized mRNAs found in the BAC screen. A-Schematic of the localization classes. The mRNAs are in red; nuclei are delimited by blue lines and cellular areas by black lines. B-Localized mRNAs. Legend as in Figure 1B. The name of the BAC tagged gene is indicated (top of each mRNA panel), as well as the localization class (italics). Scale bar: 10 microns.

**Figure 4.**
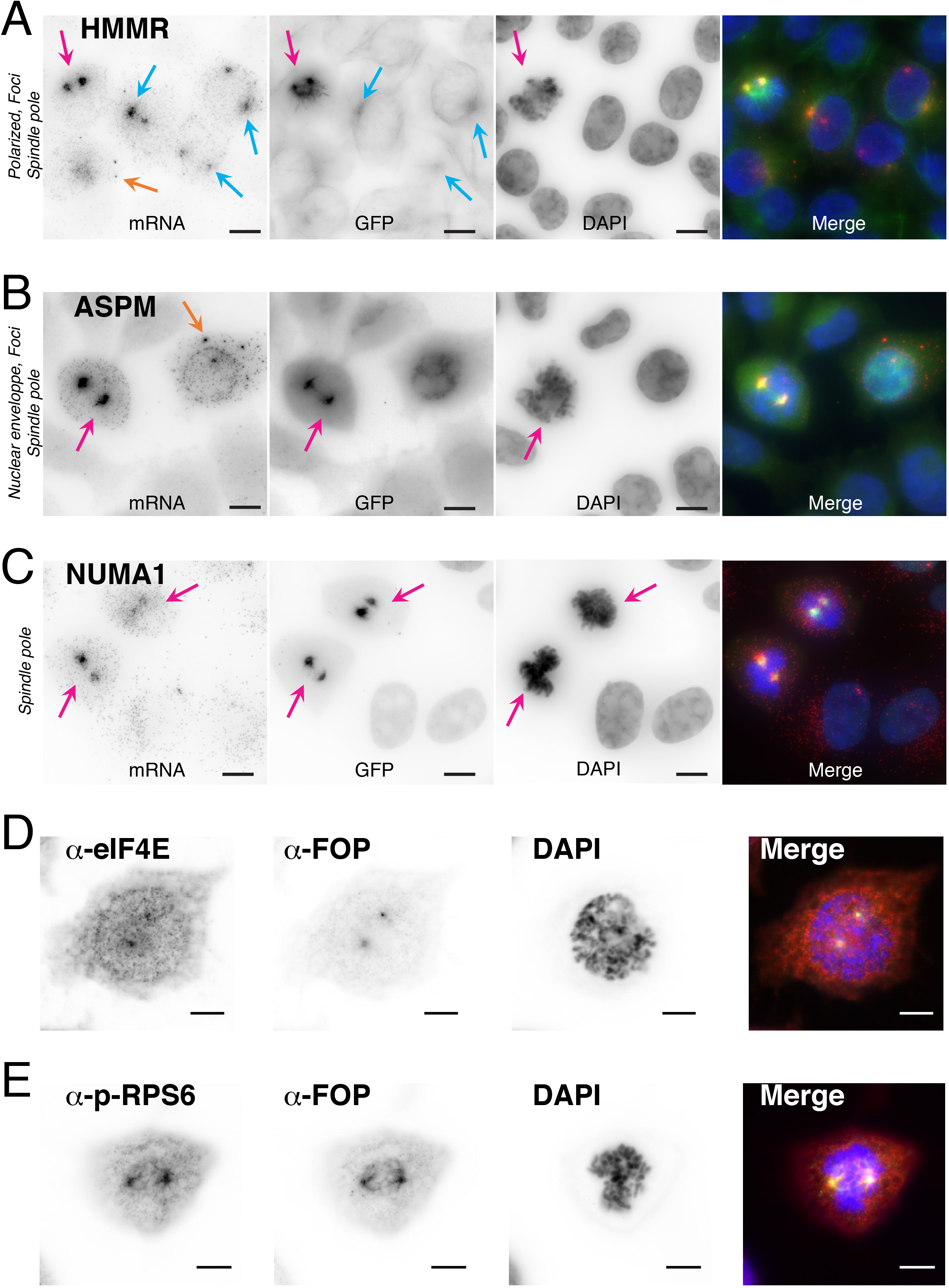
Localization of mRNAs and translation factors on centrosomes. A-C-Localization of tagged transcripts coding for HMMR (A), ASPM (B), and NUMA1 (C). Legend as in Figure 1B. The localization class is indicated on the left. Orange arrows point to mRNA foci; pink arrows point to mitotic centrosomes of cells in prophase; light blue arrows point to centrosomes of cells in interphase. Scale bar: 10 microns. D-E-Localization of eIF4E (D) and phopsho-RPS6 (E) in early mitotic cells. The α-FOP antibody (green and black) labels centrosomes. The staining of translation factors is in red and black. Scale bar: 10 microns.

**Table 1.** Summary of the localized mRNAs found in the screen.

| Localization pattern | Compartment where RNA localize | Gene | Colocalization with the encoded protein | Puromycin sensitivity of RNA localization |
| --- | --- | --- | --- | --- |
| Protrusion | Protrusion | KIF1C, KIF4A, KIF5B, MYH3, DYNLL2 | KIF1C | KIF4A<br>KIF5B and DYNLL2: nd |
| Foci | P-body | See Table S2 (15 species) | CRKL | nd |
| Foci | Translation factories | DYNC1H1, ASPM, BUB1, CTNNB1/β-catenin | -CTNNB1<br>-DYNC1H1, ASPM, BUB1 (nascent). | All |
| Foci | Centrosome | HMMR | yes | yes |
| Mitotic | Spindle poles | ASPM, NUMA1, HMMR | All (during mitosis) | ASPM, HMMR<br>NUMA1: nd. |
| Nuclear edge | Nuclear envelope | ASPM, SPEN | ASPM (nascent)<br>SPEN: nd | All |
| Perinuclear | Golgi | AKAP9 | yes | yes |
| Perinuclear | Endosome | AP1S2 | yes | yes |
| Cell edge | Cell edge | FLNA | yes | nd |
| Perinuclear | Mitochondria | AKAP1 | yes | yes |
| Perinuclear | RE | HSP90B1, ATP6A2 | yes | yes, ATP6A2: nd |
| Intranuclear | Intranuclear | See Table S2 (11 species) | nd | nd |

Most mRNAs displayed a single localization pattern, however, some showed several (Table 1 and S2). For instance, ASPM mRNAs simultaneously localized at the nuclear envelope and in foci of interphase cells, while it localized at the mitotic apparatus during cell division (Figure 4B). Likewise, HMMR mRNAs localized in both P-bodies and at centrosomes (Figure 4A and S4A). These examples highlight the complexity and dynamic nature of RNA localization even in a simple cell culture system.

### Supervised and unsupervised machine learning precisely quantify RNA localization and highlights the heterogeneity of this process

We recently developed a pipeline to analyze RNA localization in a quantitative and unbiased manner (Samacoits et al., 2018; Figure 5A). Here, we further extended and improved the pipeline. In particular, unlike our previous approach, we now explicitly allow for pattern mixtures, i.e. each cell can belong to several categories. Our method first segments cells and nuclei, detects single RNA molecules, and then calculates a series of features that describe the spatial properties of the RNA point cloud, both with respect to inter-point distance distributions and the distance to landmarks, here the nuclear and cytoplasmic membranes (Figure 5A; see details in STAR Methods). These features are mostly invariant to RNA concentration and they allow to use unsupervised or supervised approaches to classify each cell in a series of patterns. For this analysis, we first generated a manually annotated dataset. We selected several localized mRNAs together with several genes annotated as random (KIF20B, MYO18A, SYNE2, PLEC), and we then manually annotated individual cells falling into the different patterns (810 cells in total). We then investigated the distributions of these annotated cells in the multi-dimensional feature space. After reducing dimensionality with a t-SNE plot (Figure 5B), cells with different patterns were situated in different regions of the embedded feature space. Moreover, cells with the same annotations clustered in the same region whether they corresponded to the same gene or not. Thus, the features captured key parameters of the localization pattern that recapitulated the manual annotations.

**Figure 5.**
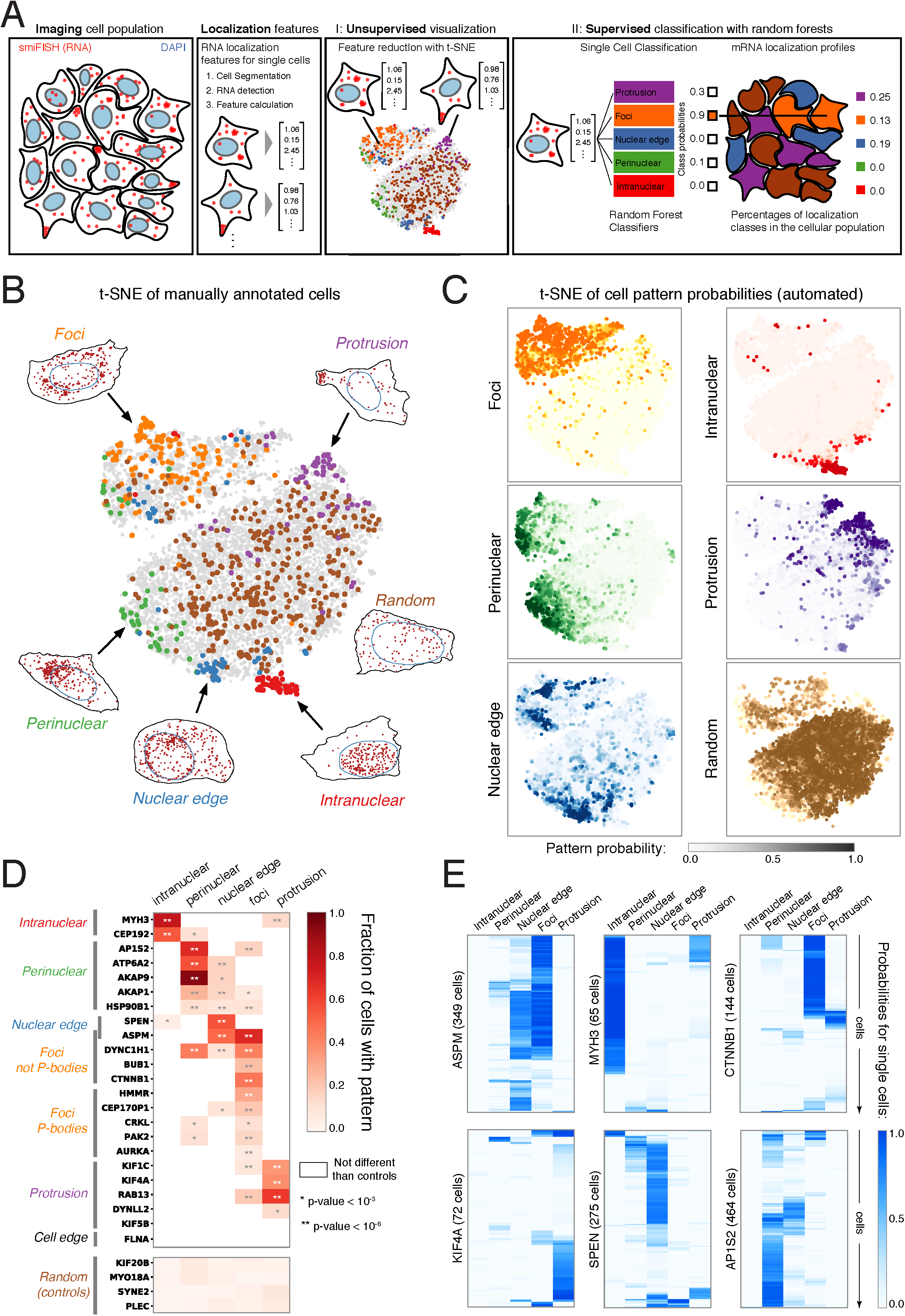
Quantitative analysis of mRNA localization by machine learning approaches. A-Schematic of the analysis pipeline. B-t-SNE plot of manually annotated cells. Each point represents a cell in the feature space after a t-SNE transformation. Cells manually annotated as ‘*Foci*’: orange; ‘*Protrusion*’: violet; ‘*Perinuclear*’: green; ‘*Nuclear edge*’: blue; ‘*Intranuclear*’: red; ‘*Random*’: brown. Cells that were not annotated or that had multiple patterns are colored in grey. One example cell is shown for each class (black: contour of the cytoplasm, red: RNA detection result). C-Visualization of Random Forest classification probabilities in the embedded feature space (t-SNE). Cells are colored according to the probability assigned by each indicated classifier; color code as in B. D-Heatmap depicting the fraction of cells classified in the indicated pattern, for the different genes analyzed by the automated pipeline. The bottom panel shows the control, randomly localized mRNAs. The top panel shows mRNAs with localization preference. Only values significantly different from negative controls (p-value <10^-3^) are colored. One star: probability between 10^-3^ and 10^-6^; two stars: probability less than 10^-6^. The manual annotations of the genes are indicated on the left with the same color codes as in B. E-Heatmaps depicting the probability of single cells to have each of the patterns, for the indicated genes. Each row corresponds to a cell, and the color indicates the probabilities of the cell to be classified in the indicated localization pattern.

Next, we developed a pipeline able to assign multiple patterns to a single cell. For this, we trained several random forest classifiers, each one recognizing one pattern (*‘protrusion’, ‘foci’, ‘nuclear edge’, ‘perinuclear’, ‘intranuclear’*), reaching an average classification accuracy between 92 to 100% (see Figure S5C). We then analyzed all the cells of the different genes (9710 cells and 27 genes). To investigate the agreement between unsupervised and supervised analysis, and to understand the biological signal corresponding to different regions of the t-SNE plot, we determined the probabilities of cells to be in a particular pattern, using each of the classifier, and plotted this probability on top of the t-SNE representation (Figure 5C). Remarkably, cells with a high probability of a given pattern concentrated in the same area of the t-SNE plot, thus providing a direct interpretation of the regions with respect to the localization pattern they represent (Figure 5C). This also indicated that the assignments of the classifiers were consistent with the manual annotations.

Finally, we analyzed RNA localization at the gene level. Individual cells were assigned to a pattern if the probability given by the random forest classifier for that pattern was higher than 0.5. The fractions of cells in each pattern were then represented in a heatmap, either directly (Figure S5E), or after filtering out cases where the proportion of cells with localized mRNA was not statistically different from that of the control random genes (Figure 5D, threshold: p-value < 10^-3^). There was a remarkable agreement between the automated and manual classification. Except for FLNA and KIF5B, all the manual annotations were confirmed with high degree of statistical significance (p-value < 10^-3^). FLNA was the only mRNA manually annotated as ‘*cell edge*’ and we decided not to study it further. KIF5B indeed localized weakly in protrusions of HeLa cells (i.e. in a small number of cells), although the localization pattern was nicely seen in neuronal cells (Figure S1B). The other protrusion mRNAs were statistically different than the controls (p-value < 10^-3^), with 14% (DYNLL2) and up to 62% (RAB13) of the cells classified in that pattern. The two genes manually annotated as ‘*intranuclear*’ had more than 55% of the cells assigned in that category. Likewise, the two ‘*nuclear edge*’ genes, ASPM and SPEN, had respectively 50% and 55% of cells in this category, while the ‘*perinuclear*’ genes (AKAP1, AKAP9, AP1S2, ATP6A2 and HSP90B1) ranged from 13% (HSP90B1), to 93% (AKAP9). The former value may seem low, but it is nevertheless highly statistically significant (p-value of 7.10^-15^). Also note that HSP90B1 mRNAs colocalized with their encoded proteins, providing further evidence that they are localized (see below). For the pattern ‘*foci*’, the non-P-body genes (ASPM, BUB1, CTNNB1 and DYNC1H1) had a high proportion of cells classified in that pattern (24% for BUB1 and up to 69% for ASPM), and an average proportion of mRNA in foci varying from 9% (BUB1) to 28% (CTNNB1; Figure S5B). In contrast, the mRNAs accumulating in P-bodies had a comparatively smaller fraction of cells classified in the ‘*foci*’ pattern (7 to 31%), as well as a smaller fraction of the mRNAs in foci (2 to 15%). Note that our estimation of the fraction of mRNA in foci correspond to a lower bound because we only counted groups of molecules where each molecule was closer than 350 nm to the group, while foci can be larger. Nevertheless, and taking into account our stringent criterion, our quantification was in the lower range of previously reported quantifications of P-body mRNAs (10-20% of the total mRNAs; Pillai et al., 2005; Hubstenberger et al., 2017).

When comparing individual cells, the quantifications indicated that RNA localization can be highly variable even for the same mRNA. For instance, the fraction of CTNNB1 mRNA in foci varied from 0 to more than 60%. To analyze this phenomenon in more details, we plotted the cells of each single gene on t-SNE plots (Figure S6), and we also generated heatmaps depicting the probabilities of the different patterns for all the individual cells (Figure 5E and S6). This provided a very detailed view of RNA localization at the level of single cells. It confirmed the intercellular variability of RNA localization as cells of a given gene were dispersed of the t-SNE plot, although an accumulation in the expected area of the plot was visible (Figure S6). Likewise, the pattern probabilities varied greatly from cell to cell (Figure S6). It was also apparent that a large number of cells had several patterns simultaneously. In particular, many DYNC1H1 cells were classified as both ‘*perinuclear*’ and ‘*foci*’, and many ASPM cells were both ‘*nuclear edge*’ and ‘*foci*’ (in agreement with the manual annotations). Likewise, MYH3 cells were frequently ‘*intranuclear*’ and ‘*protrusion*’. In conclusion, this quantitative analysis corroborated the manual annotations and revealed a high degree of heterogeneity of RNA localization across different cells. This heterogeneity is a general phenomenon since it is seen with nearly all the genes analyzed here.

### Identification of potential cases of locally-translated mRNAs

Having analyzed the localization patterns by a quantitative and unbiased approach, we then examined the colocalization of the mRNAs with their encoded protein. Although accumulation of mRNAs in foci was the most frequent pattern (19/32; see Table 1), only two mRNAs displayed a faint but detectable GFP signal in their foci: CRKL and CTNNB1/β-catenin (see Figure 3B and S3 for CRKL, and Figure S10A for CTNNB1/β-catenin). For the other localization patterns, 9 mRNAs colocalized with their protein (2% of the screened mRNAs; Table 1): AKAP1, AKAP9, AP1S2, ASPM, ATP6A2, FLNA, HMMR, NUMA1 and HSP90B1 (Figure 3B and 4). In some cases, the mRNA/protein colocalization was expected. For instance, both HSP90B1 and ATP6A2 contain a signal peptide leading to translation on the ER, and HSP90B1 is a resident ER protein while ATP6A2 localizes in endo-lysosomes close to this compartment (Figure 3B). Similarly, AKAP1 encodes an RNA-binding protein localizing to the surface of mitochondria, and this transcript thus belongs to the known class of mitochondrion-localized mRNAs (Sylvestre et al., 2003). The other proteins localized to cellular structures that were not previously known to use local translation as a targeting mechanism. This included the clathrin adaptor AP1S2 mRNA that localized on endosomes and the AKAP9 mRNA that accumulated at the Golgi. Below, we explore these cases in more detail together with mRNAs localizing around centrosomes (ASPM, NUMA1 and HMMR), at the nuclear envelope (ASPM and SPEN), or in cytoplasmic foci that were not P-bodies (ASPM, BUB1, DYNC1H1, and CTNNB1/β-catenin).

### ASPM, NUMA1 and HMMR mRNAs localize to centrosomes in a cell-cycle dependent manner

The HMMR protein is known to localize on microtubules and centrosomes and to have spindle-promoting activities (Groen et al., 2004; Maxwell et al., 2003). The HMMR mRNAs accumulated in the peri-centrosomal region in interphase and much more strongly so in mitosis, and it colocalized at the centrosome with its protein (Figure 4A and S7A). Moreover, using a RPE1 cell line stably expressing a Centrin2-GFP protein, we confirmed that the endogenous HMMR mRNAs preferentially localized near centrosomes (Figure S7B).

The other mRNAs localizing at centrosomes were ASPM and NUMA1. Both proteins control spindle function during mitosis (Kouprina et al., 2005; Radulescu and Cleveland, 2010). Interestingly, these two proteins had similar localization patterns. In interphase, both localized to the nucleoplasm while during mitosis, they concentrated on the spindle poles and weakly stained microtubules (Figure 4B-C; pink arrow points to cells in prophase). Remarkably, the localization of ASPM and NUMA1 mRNA displayed a similar dynamic during the cell cycle (Figure 4B-C). In interphase, these mRNAs were dispersed throughout the cytoplasm, with ASPM mRNAs additionally localized in foci and decorated the nuclear edge. However, both mRNAs re-localized to centrosomes during mitosis, where they became highly concentrated and colocalized with their GFP-tagged proteins (Figure 4B-C, pink arrow indicates cells in early mitosis). Identical localization patterns were observed for the endogenous mRNAs (Figure S7C-D).

The co-localization of HMMR, ASPM and NUMA1 mRNAs with their proteins suggested that they were translated locally at centrosomes. To confirm this possibility, we analyzed the localization of two translation factors: eIF4E and the phosphorylated form of the ribosomal protein RPS6 (p-RPS6). Immuno-fluorescence showed that the endogenous eIF4E and p-RPS6 proteins were present throughout the cells but also accumulated at mitotic centrosomes (Figure 4D-E, see cells in prophase). Therefore, not only mRNAs but also the translational apparatus was present at the spindle poles. This suggests the presence of a specific translational program occurring on mitotic centrosomes.

### The localization of ASPM and SPEN mRNAs at the nuclear envelope is translation-dependent

ASPM-GFP mRNAs accumulated at the nuclear envelope during interphase (Figure 4B; quantification in Figure 5E). In addition, SPEN-GFP mRNAs, which encode a nuclear protein, were also enriched around the nuclear envelope (Figure 3B; Figure 5E). Labelling of the nuclear pores using either a CRM1-GFP fusion or an antibody against NUP133 (Figure S8A-B), showed that ASPM and SPEN mRNAs localized close to nuclear pores, rather than between them. This localization could result from two mechanisms. First, mRNAs could transiently localize at the pores on their way out of the nucleus. Second, they could localize at the cytoplasmic side of the pore to facilitate re-entry of newly translated proteins. To distinguish between these possibilities, we inhibited either transcription with actinomycin D, or translation with puromycin. After 1h of actinomycin D treatment, SPEN and ASPM mRNAs still localized at the nuclear envelope (left panels in Figure 6A; Figure S8C and 6B for quantifications). On the contrary, both mRNAs became dispersed after 1h exposure to puromycin (Figure 6A, S8C and S9A for quantifications). We then used cycloheximide, which inhibits translation by freezing the ribosomes on the mRNAs, in contrast to puromycin that induces premature termination and releases the nascent peptide. Cycloheximide had no effects on the localization of ASPM mRNAs (Figure 6A left panels and 6B), indicating that their localization did not require protein synthesis per se, but more likely the presence of nascent proteins on polysomes. Therefore, ASPM and SPEN mRNAs at the nuclear envelope are not in transit to the cytoplasm but localize on the nuclear envelope by a translation-dependent mechanism requiring the nascent protein.

**Figure 6.**
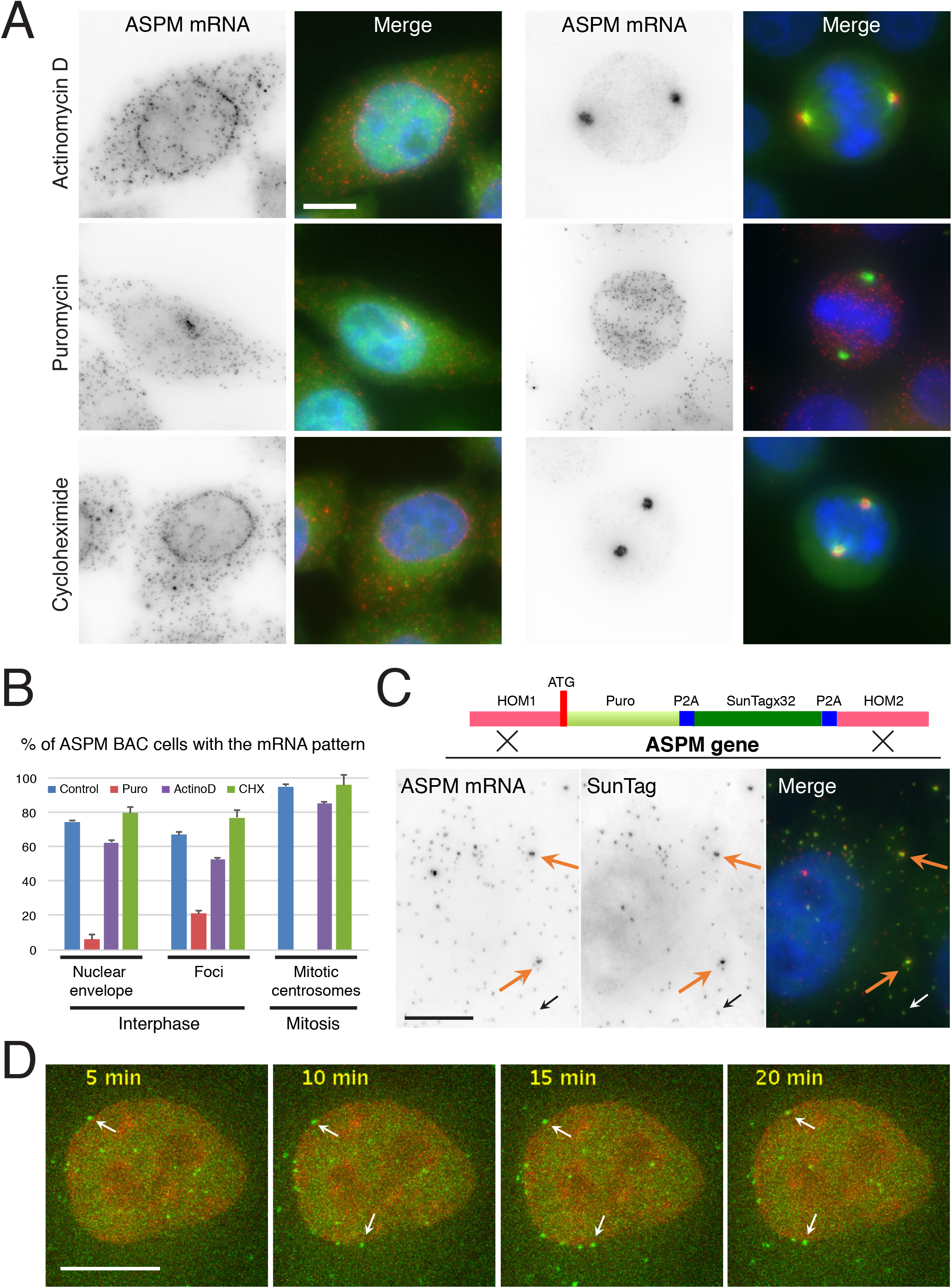
Localization of ASPM mRNAs requires active translation. A-Localization of ASPM mRNA to the nuclear envelope and mitotic centrosomes is abolished by puromycin. Panels depict micrographs of HeLa cells expressing the tagged ASPM BAC, after 1h of treatment with the drugs indicated on the left. The RNA foci in the puromycin panel is a transcription site. Scale bar: 10 microns. B-Quantification of the localization of the tagged ASPM mRNAs. The graph depicts the percent of cells expressing the tagged ASPM mRNAs and having the transcript localized in the indicated pattern (using manual annotations). C-Foci of ASPM mRNAs contain polysomes. Top: schematic depicting the insertion of the SunTag at the N-terminus of ASPM protein. Bottom: micrograph of HeLa cells with a SunTagged ASPM allele and showing ASPM mRNA (by smiFISH, left and red), and the signal from the SunTag (middle and green). Blue: DAPI staining. White and black arrows: a single mRNA positive for the SunTag; orange arrow: an mRNA foci positive for the SunTag. Scale bar: 10 microns. D-ASPM mRNAs can be translated at the nuclear envelope. Legend as in C, except that cells are imaged live and white arrow points to polysomes anchored at the nuclear envelope. The DNA (red) is stained with SiR-DNA.

### ASPM mRNAs are translated at the nuclear envelope

We and others recently demonstrated that the SunTag can be used to visualize translation of single mRNPs in live cells (Pichon et al., 2016; Wang et al., 2016; Wu et al., 2016; Yan et al., 2016; reviewed in Pichon et al., 2018). The SunTag is a repeated epitope and in cells expressing an scFV-GFP monochain antibody directed against this epitope, the antibody binds the repeated tag as soon as it emerges from the ribosome, allowing the visualization of single molecules of nascent proteins (e.g. monosome and polysomes). To confirm that ASPM mRNAs were translated at the nuclear envelope, we tagged the endogenous gene using CRISPR genome editing. Homologous recombination allowed to obtain heterozygous clones where 32 SunTag repeats were fused at the N-terminus of ASPM proteins (Figure S8D). The endogenous ASPM transcripts were then labelled by smiFISH, revealing both tagged and untagged mRNAs. This showed that bright spots of scFv-GFP colocalized with single ASPM mRNAs (Figure 6C; white arrows). These spots disappeared after a 20 minutes puromycin treatment, confirming that they were polysomes (Figure S8E). We then performed time-lapse analyses of live cells, recording one image stack every 40 seconds for 50 minutes (see Movie and still images in Figure 6D). Remarkably, some polysomes localized at the nuclear envelope. While cytoplasmic polysomes moved too rapidly to be tracked, the ones at the envelope remained immobile for extended periods of times (29 minutes in average). Thus, a fraction of ASPM mRNAs was stably anchored and translated at the nuclear envelope.

### Translation inhibition frequently prevents mRNA localization

Since the localization of ASPM mRNA at the envelope is translation dependent, we tested whether this was also the case for its localization at centrosomes. Puromycin abolished its localization, while actinomycin D and cycloheximide had little effect (Figure 6A, right panels; Figure 6B for quantification). The effect of puromycin was then tested on the other localized mRNAs. After 1 hour of treatment, KIF1C and MYH3 mRNAs still localized to cytoplasmic protrusions (Figure 7A and 7B for quantifications). In contrast, HSP90B1, HMMR, AP1S2, AKAP1, KIF4A and AKAP9 mRNAs all became delocalized and lost co-localization with their encoded protein when translation was inhibited (Figure 7C-E; Figure S9A). This data demonstrated that translation is required for the localization of these mRNAs.

**Figure 7.**
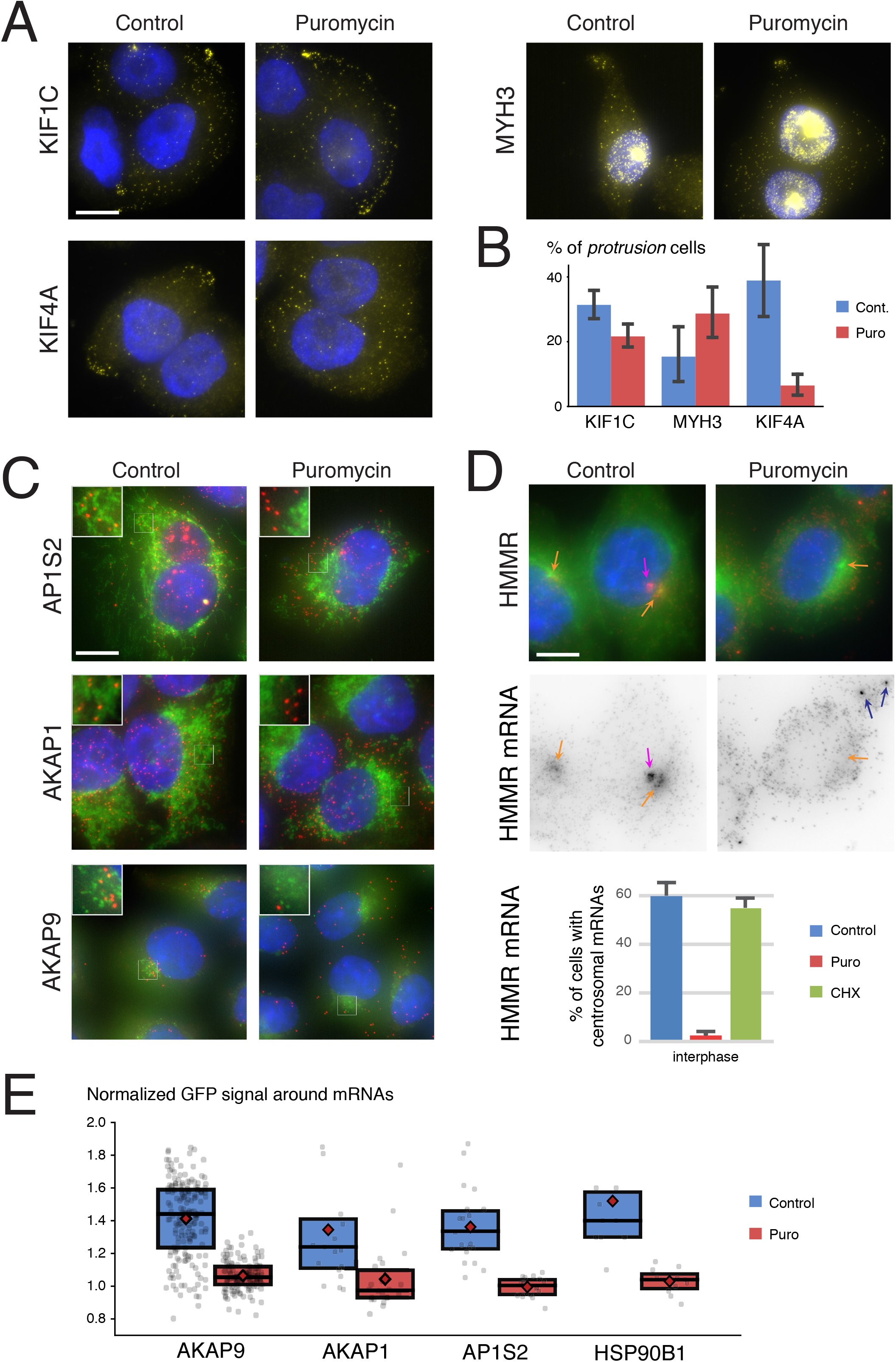
Transcript localization is sensitive to translation inhibition. A-Transcripts localizing in cytoplasmic protrusions are diversely affected by puromycin. Images are micrographs of HeLa cells expressing the indicated BAC-tagged gene, treated or not with puromycin for 1 hour, and hybridized with Cy3-labelled probes against the GFP-IRES-Neo tag (yellow). Blue: DAPI staining. Scale bar: 10 microns. B-Fraction of cells classified in the pattern ‘*protrusions*’ by the automated classifier. The graph depicts the percent of cells displaying mRNA accumulation at the cell periphery, either in absence of treatment (blue bars), or after 1 hour of incubation with puromycin (orange bars). Error bars represent the 95% confidence interval computed by bootstrapping. C-Puromycin treatment disrupts the localization of AP1S2, AKAP1 and AKAP9 mRNAs. Legend as in Figure 1B; puromycin-treated cells were incubated 1 hour with the drug. Insets: zoom of the boxed areas. Scale bar: 10 microns. D-Puromycin, but not cycloheximide, disrupts the localization of HMMR mRNA to centrosomes. Legend as in Figure 1B, puromycin treatment was for 1 hour. Scale bar: 10 microns. Dark blue arrows: mRNA in foci (P-bodies, see Figure S6). Orange arrows: centrosomes; pink arrow: transcription site. Bottom: the graph depicts the percent of cells expressing the tagged HMMR mRNA and accumulating it at the centrosome during interphase (using manual annotations). E-Quantification of the co-localization between mRNAs and their encoded protein. The graphs display the normalized GFP fluorescence intensities around the indicated BAC-tagged mRNAs. Values are averaged for each cell, and each dot is a cell. The box is the second and third quartiles; the black bar is the median and the red diamond is the mean. The puromycin samples are statistically different from the untreated ones (p-value < 0.05 with a two-sample Kolmogorov-Smirnov test).

### Non P-bodies mRNA foci correspond to specialized translation factories

Four mRNAs accumulated in foci that were distinct from P-bodies (BUB1, DYNC1H1, CTNNB1/β-catenin and ASPM; Figure S5A). The dynein heavy chain DYNC1H1 and the ASPM protein were described above. BUB1 is a checkpoint kinase that verifies the attachment of microtubules to kinetochores at the onset of mitosis (Saurin, 2018), and β-catenin is the key transcription factor of the Wnt signaling pathway (Grainger and Willert, 2018). SmiFISH confirmed that the four endogenous mRNAs accumulated in foci (Figure S1A; S7C; S9E and see below S10F). Moreover, dual-color smiFISH showed that these foci did not co-localize together, indicating that they were distinct structures (Figure 8A). We then inhibited translation with puromycin, using an mRNA accumulating in P-bodies as control (AURKA). After 1h of treatment, the non-P-body foci virtually disappeared (Figure 8B-C), while the accumulation of AURKA mRNA in P-bodies actually increased. Thus, translation inhibition specifically disrupted the mRNA foci that were not P-bodies. We previously used the SunTag to show that DYNC1H1 is translated in the mRNA foci (Pichon et al., 2016). Using the CRISPR SunTag clone described above, we found that ASPM mRNAs were similarly translated in the mRNA foci (Figure 6C, orange arrow; 70% of the foci have a SunTag signal). A 32xSunTag repeat was then introduced at the N-terminus of the BUB1 protein using CRISPR gene editing. BUB1 mRNAs were detected by smiFISH and were found to be frequently translated in the foci (Figure 8D-E; 70% of the foci contained a SunTag signal), in contrast to mRNAs located outside foci that were rarely translated (1% of the time). Taken together, this data demonstrated that ASPM, BUB1 and DYNC1H1 are translated in mRNA foci, which thus correspond to specialized translation factories.

**Figure 8.**
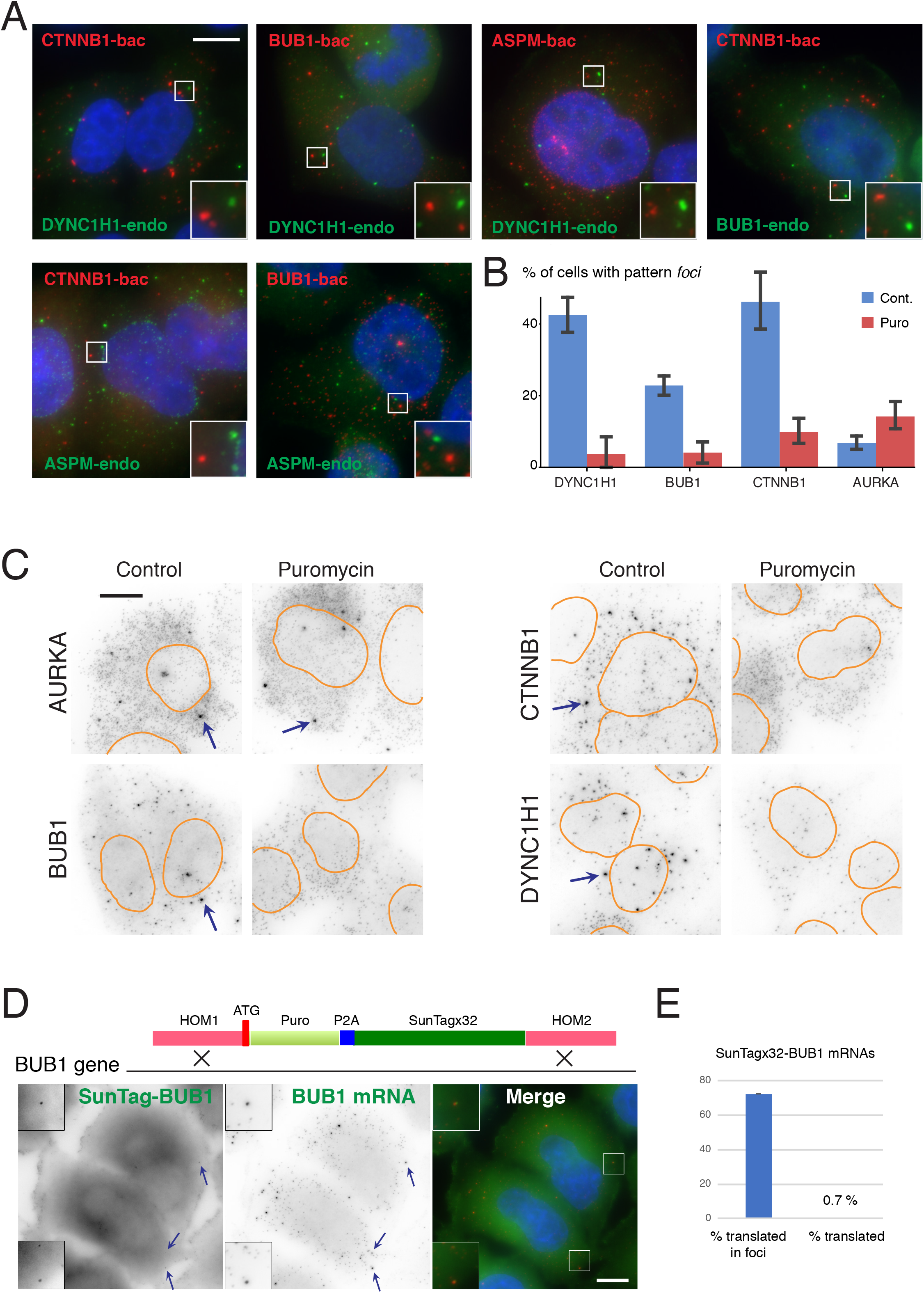
Non-P-body mRNA foci are translation factories. A-Foci containing BUB1, ASPM, CTNNB1 and DYNC1H1 mRNAs do not co-localize with each other. Panels are micrographs of HeLa cells expressing the indicated BAC-tagged gene and hybridized with Cy3-labelled probes against the GFP-IRES-Neo sequence (red), together with Cy5-labelled probes against the indicated endogenous mRNA (green). Blue: DAPI staining; scale bar: 10 microns. B-C-Puromycin dissociates non-P-body foci. B-Fraction of cells classified in the pattern ‘*foci*’ by the automated classifier, for the indicated genes and in presence (red) or absence (blue) of puromycin for 1h. Error bars represent the 95% confidence interval computed by bootstrapping. C-Micrographs are images of HeLa cells expressing the indicated BAC and hybridized with Cy3-labelled probes against the GFP-IRES-Neo tag. Arrows point to mRNA foci, and the nuclear outline is shown in orange. Scale bar: 10 microns. D-BUB1 mRNAs are translated in mRNA foci. Top: schematic depicting the insertion of the SunTag at the N-terminus of the BUB1 protein. Bottom: micrograph of HeLa cells with a SunTagged BUB1 allele and showing BUB1 mRNA (by smiFISH; middle and red), and the signal from the SunTag (left and green). Blue: DAPI staining. Arrows: mRNA foci that are positive for the SunTag. Insets: zoom of the boxed areas; scale bar: 10 microns. E-Quantification of translated SunTagged BUB1 mRNA. The graph represents the percent of BUB1 mRNA foci positive for the SunTag (left bar), and the percent of single BUB1 mRNAs also positive for the SunTag (right bar), in cells with one BUB1 allele tagged with the SunTag. Error bars: standard deviation.

### CTNNBl/β-catenin mRNA foci are sites of co-translational protein degradation

We then analyzed in more details CTNNB1/β-catenin. This protein has two roles: it bridges E-cadherin to the actin cytoskeleton at adherens junctions, and it acts as the main transcription factor of the Wnt pathway (for review, see Grainger and Willert, 2018; MacDonald et al., 2009). The Wnt signaling pathway is essential during development, is often a key actor during tumorigenesis, and involves a fast activation of β-catenin expression by Wnt. The β-catenin protein localized at adherens junction is stable whereas the one synthesized in the cytoplasm is rapidly degraded (reviewed in Stamos and Weis, 2013). However, the cytoplasmic protein is stabilized in presence of a Wnt signal, and can then accumulate in the nucleus to activate transcription of target genes. In the β-catenin BAC cells, the GFP-tagged protein was weakly expressed. It accumulated at sites of cell-cell contacts as expected, and also showed a weak staining in the brightest RNA foci (Figure S10A). Because the maturation of the GFP chromophore is slow and therefore inadequate to map translation sites, we combined smFISH with immunofluorescence, using an antibody binding the N-terminus of β-catenin. The mRNA foci contained high levels of β-catenin N-termini (Figure S10B). Since these foci were also dissolved when translation was inhibited (see Figure 8B-C), these data indicated that β-catenin was translated in the mRNA foci.

Interestingly, there was a small fraction of the BAC cells where β-catenin mRNAs did not form foci, and instead were dispersed as single molecules in the cytoplasm. Furthermore, these cells expressed high levels of nuclear β-catenin-GFP, suggesting that the Wnt pathway had been activated and was responsible for the disappearance of the foci (Figure S10C). To test this hypothesis, we activated the Wnt pathway by incubating the BAC cell line with the WNT3A protein. Indeed, the mRNA foci disappeared after 30 minutes, concomitant with a higher expression of β-catenin-GFP and its accumulation in the nucleus (Figure 9A and 9E for quantifications). This observation established a link between the presence of mRNA foci and the degradation of the β-catenin protein. One hypothesis to explain these results would be that β-catenin degradation takes place co-translationally in the foci. Degradation of β-catenin requires Axin, APC and the kinases CK1α and GSK3, which altogether form the “destruction complex” (Stamos and Weis, 2013). In absence of Wnt, this complex binds β-catenin and targets it for degradation. In presence of Wnt, its components become recruited to the plasma membrane and they can no longer interact with β-catenin (Grainger and Willert 2018, Stamos and Weis, 2013, MacDonald et al., 2009). Axin is an essential player of the destruction complex and it acts as a scaffolding protein (Grainger and Willert 2018, Stamos and Weis, 2013, MacDonald et al., 2009). It binds β-catenin as well as APC, CK1α and GSK3, and its role is to bring β-catenin in proximity to the kinases CK1α and GSK3 (3-5). β-catenin is first phosphorylated on residue S45 by CK1α, leading to its recognition by GSK3, which phosphorylates it on residues T41, S37 and S33. Phosphorylated β-catenin is then recognized by the E3 ubiquitin ligase β-TrCP, which targets it for proteasomal degradation. APC is a very large protein that is essential for β-catenin degradation but whose exact molecular function is unclear. It occurs as a dimer and each monomer has more than 30 tandemly repeated binding sites for β-catenin.

**Figure 9.**
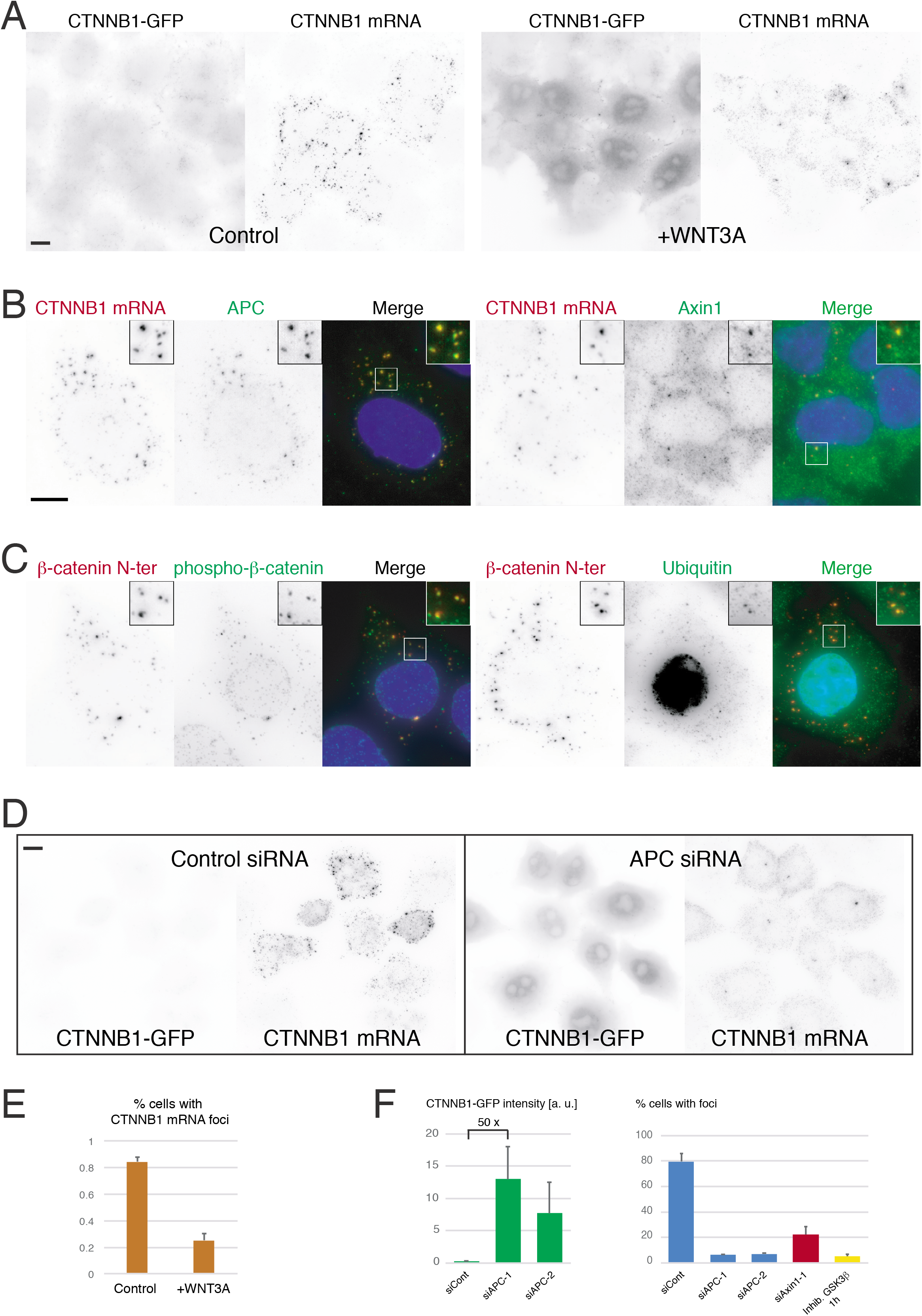
CTNNB1 mRNA foci correspond to regulated sites of protein degradation. A-WNT3A signaling induces β-catenin (CTNNB1) protein expression and dissolve mRNA foci. Images are micrographs of HeLa cells expressing the tagged CTNNB1 BAC, and hybridized *in situ* with Cy3-labelled probes against the tag (panels CTNNB1 mRNA), and treated or not with WNT3A conditioned media for 30 minutes (+WNT3A and control panels, respectively). CTNNB1-GFP panels: signal from the GFP channel, with the grey levels scaled identically for the WNT3A and control. Scale bar: 10 microns. B-β-catenin mRNAs foci contain APC and Axin. Images are micrographs of HeLa cells expressing the tagged CTNNB1 BAC and both hybridized in situ with Cy3-labelled probes against the tag (panel CTNNB1 mRNA and red in the merge), and incubated with Cy5-labelled anti-APC (left; panel APC and green in the merge) or anti-Axin1 antibodies (right; panel Axin1 and green in the merge). Blue: DAPI staining. Scale bar: 10 microns. C-β-catenin mRNAs foci contain phosphorylated β-catenin and ubiquitin. Images are micrographs of HeLa cells expressing the tagged CTNNB1 BAC and stained by immuno-fluorescence with antibodies against the N-terminus of β-catenin (panel β-catenin N-ter and red in the merge), and with antibodies against phosphorylated forms of β-catenin (left; panel phospho-β-catenin and green in the merge) and ubiquitin (right; panel Ubiquitin and green in the merge). Blue: DAPI staining. Scale bar: 10 microns. D-Knock-downs of APC dissolve β-catenin mRNAs foci and induce β-catenin protein expression. Images are micrographs of HeLa cells expressing the tagged CTNNB1 BAC, and hybridized *in situ* with Cy3-labelled probes against the tag (panels CTNNB1 mRNA), and treated or not with APC siRNAs, as indicated. CTNNB1-GFP panels: signal from the GFP channel, with grey levels scaled identically for the APC siRNA and control. Scale bar: 10 microns. E-Quantification of the effect of WTN3A. Graph represent the percent of cells expressing β-catenin mRNA and displaying mRNA foci (using manual annotations), in control cells or cells treated with WNT3A-conditionned medium for 30 minutes. Error bars represent the standard deviation of replicates. F-Quantification of the effect of APC and Axin1 knock-down, and GSK3 inhibition. The left graph depicts the intensity levels of β-catenin-GFP protein in the nucleus, and the right graph displays the percent of cells expressing β-catenin mRNA and having mRNA foci (using manual annotations), for the indicated condition. Error bars: standard deviation.

Labelling with anti-APC or anti-Axin1 antibodies revealed that the β-catenin mRNA foci contained high levels of APC and were also enriched for Axin1 (Figure 9B). Moreover, Axin1 could be detected in polysomal fractions in a translation-dependent manner (Figure S10D), suggesting that it binds β-catenin co-translationally. We then immuno-localized phosphorylated β-catenin (on residues S33 and S37), to test its presence in the mRNA foci. In this case, we used N-terminal labeling of β-catenin as a proxy for the mRNA foci because the anti-phospho antibodies were not compatible with smFISH (Figure 9C). Indeed, phospho-β-catenin accumulated in the mRNA foci. In addition, we could also detect cross-linked chains of ubiquitin itself (Figure 9C). Altogether these data indicated that the mRNA foci were sites of both β-catenin synthesis and degradation, i.e. sites of co-translational protein degradation. Foci formation was further regulated by Wnt signaling, thereby allowing a fast post-translational response.

Interestingly, Axin can oligomerize and APC can cross-link these oligomers to make large aggregates (Grainger and Willert 2018, Stamos and Weis, 2013, MacDonald et al., 2009), suggesting that these factors could be involved in foci formation. Indeed, when APC was knocked-down with siRNA, we observed a 50-fold increase of β-catenin levels and, remarkably, a disappearance of the mRNA foci (Figure 9D and 9F). Likewise, mRNA foci were disrupted upon Axin1 knock-down. In contrast, when we increased Axin1 levels by inhibiting Tankyrase with the small molecule LG-007 (Huang et al., 2009), the endogenous β-catenin mRNA formed more foci in HEK293 cells: foci were visible in 16% of the cases in untreated cells, and this increased to 32% after LG-007 treatment for 2h (Figure S10F-G). Finally, we also tested a pharmacological inhibition of GSK3. Indeed, APC and Axin1 are themselves phosphorylated by GSK3, and this promotes complex formation and their interaction with β-catenin (Ha et al., 2004; Wu and Pan, 2010). Remarkably, GSK3 inhibition for 1h nearly completely dissolved β-catenin mRNA foci. These data suggest that foci formation relies on the co-translational recognition of β-catenin by APC and Axin1, and the multimerization of these factors.

## Discussion

In this study, we performed a dual RNA/protein localization screen and analyzed more than 500 genes. We found 32 localized mRNAs, with 15 accumulating in P-bodies and the others being locally translated (Table 1). We also discovered a number of unexpected features, and in particular that: (i) co-translational mRNA targeting is frequent and occurs at unexpected locations; and (ii) mRNAs can be translated in specialized translation factories, which can perform key functions on the metabolism of nascent proteins.

### Localization of mRNA shows a high degree of heterogeneity in cells lines

We performed a quantitative analysis of mRNA localization using both unsupervised and supervised strategies. This analysis corroborated the manual annotations by confirming mRNA localization, and it further provided a detailed and quantitative view of this process at the level of single cells for many genes. This revealed that only a fraction of cells displayed the expected localization pattern, with levels ranging from 7% for AURKA to 93% for AKAP9. In addition, even in cells displaying a localization pattern, only a fraction of the molecules localized. For instance, an average of 13% of mRNA molecules accumulated in cytoplasmic protrusion, while an average of 11% concentrated in cytoplasmic foci, with large cell-to-cell variations. This indicates both inter- and intra-cellular levels of heterogeneity: genetically identical cells can localize the same mRNA in quite different ways, and within a cell, all the molecules do not necessarily share the same localization preference. This situation is quite different from what has been observed in embryos, where the RNA localization patterns appear much more stereotyped. Possibly, this reflects the diversity and heterogeneity occurring in cell lines, which has been observed in a number of ways including at the level of gene expression. It remains to be seen whether the heterogeneity of RNA localization is due to biological noise or has some functional consequences. Our data with β-catenin suggest that this heterogeneity could be important and may be linked to the physiological status of the cells (see below).

### Co-translational RNA targeting is widespread

We found a number of mRNAs that co-localize with their encoded proteins, suggesting that these are translated locally. Besides the well-known cases of proteins targeted to the ER or the mitochondria (HSP90B1, ATP6A2, AKAP1), we found that this occurred in other locations such as cytoplasmic protrusions (KIF1C), centrosomes (NUMA1, HMMR, ASPM), endosomes (AP1S2), the Golgi apparatus (AKAP9) and the nuclear envelope (ASPM and also likely SPEN). Widespread local translation may reflect the need of some proteins to be incorporated in their relevant macromolecular complex in a co-translational manner, directly at their final location. Translation at the nuclear envelope is more surprising and it could have several purposes. First, folding constraints may render import of the mature protein inefficient, for instance if the protein tends to aggregate in absence of its nuclear partners. Second, it could prevent the protein from diffusing in the cytosol and interfering with cytoplasmic processes.

One remarkable outcome of this study is that in contrast to our expectations, mRNA localization frequently requires their translation. Indeed, if one excludes mRNAs localized to P-bodies, we found that RNA-driven, translation-independent localization seems the exception rather than the rule, with only two mRNAs out of 13 falling in this category (KIF1C and MYH3, both localizing in protrusions). Moreover, the differential effects of puromycin and cycloheximide indicate that the nascent peptides appear important for mRNA targeting (see Figure 6). This is reminiscent of the SRP-dependent targeting of proteins to the ER, where the nascent peptide recruits SRP, which halts ribosomes until the entire complex docks on the SRP-receptor on the ER (Aviram and Schuldiner, 2017). In the future, it will be interesting to determine: (i) whether co-translational targeting to these other cellular locations also involves ribosome stalling, (ii) whether the mRNA contains targeting signals in addition to the nascent proteins, and (iii) whether the polysomal complex reaches destination via diffusion or active transport.

Overall, these data indicate that co-translational RNA targeting is more widespread than previously appreciated and occurs at diverse intracellular locations. This suggests that the nascent protein chain plays an important role in the targeting mechanism, and that cell may have dedicated systems to transport polysomes.

### A specific translational program occurs at mitotic centrosomes

Localization of mRNAs at mitotic centrosomes has been first observed in mollusc embryos, as a way to asymmetrically segregate mRNAs involved in developmental patterning (Lambert and Nagy, 2002). Centrosomal mRNAs have later been found during Drosophila development (Lécuyer et al., 2007), and also very recently in Zebrafish embryos, in the case of the pericentrin mRNAs that encode a centrosomal protein (Sepulveda et al., 2018). Here, we found in HeLa cells three mRNAs that localize to mitotic centrosomes together with their proteins: HMMR, NUMA1 and ASPM. We further show that the translation apparatus also accumulates there, providing a strong support to the idea that these proteins are translated directly at the spindle poles during mitosis.

The spatio-temporal localization of HMMR, ASPM and NUMA1 mRNAs is remarkably complex, with multiple patterns and localization at centrosomes occurring only at specific stages. It is interesting to note that the centrosomal localization of these mRNAs is highest in prophase (Figure 4), as in the case of pericentrin mRNAs (Sepulveda et al., 2018). Since inhibiting translation blocks mitosis entry (Epifanova et al., 1969), it is tempting to speculate that there is a specific translational program that takes place at centrosomes at the onset of mitosis, which would be required for mitosis entry.

### P-bodies and translation factories

The most frequently observed localization class was “foci”, in which mRNAs form cytoplasmic aggregates containing multiple molecules. We identified 19 such mRNAs, and 15 of the corresponding foci were P-bodies, confirming that this is an abundant localization class (Hubstenberger et al., 2017). Four mRNAs (ASPM, BUB1, DYNC1H1, and CTNNB1/β-catenin) formed foci distinct from P-bodies and also distinct from each other, and we showed that they correspond to specialized translation factories. Except for β-catenin (see below), the function of these factories is not known. However, since the mature protein does not accumulate there, they likely relate to the metabolism of the nascent protein. They could potentially be involved in the co-translational assembly of protein complexes (Pichon et al., 2016; Shiber et al., 2018; Panasenko et al., 2019), or help to locally concentrate specific chaperones or modification enzymes.

In the case of β-catenin, we showed that the factories dissolve upon Wnt signaling, concomitant to a large increase in β-catenin expression. In absence of Wnt, β-catenin is rapidly degraded and this requires APC, Axin, GSK3 and CK1α (Stamos and Weis, 2013). These factors associate with each other to form “destruction complex”, and they function by binding β-catenin and driving its phosphorylation and subsequent degradation by the proteasome (Grainger and Willert, 2018; MacDonald et al., 2009). Remarkably, the β-catenin translation factories contain APC, Axin1, phosphorylated β-catenin and ubiquitin, demonstrating that they are sites of β-catenin degradation. The high local concentrations of APC and Axin within factories may favor interactions such that all newly-made β-catenin interacts with the destruction complex, thereby preventing leakage of the system. Our discovery of the β-catenin translation factories provides a function (i.e. to aggregate polysomes) for the previously noted ability of the destruction complex to make large molecular assemblies (Mendoza-Topaz et al., 2011). This property would otherwise be difficult to rationalize, because concentrating the degradation factors in a mega-complex while the protein is made elsewhere would decrease the chances of binding. Taken together, these data revealed that co-translational protein degradation is a gene regulation mechanism enabled by translation factories.

We show that APC, Axin1 and GSK3 kinase activity are required to form the β-catenin mRNA foci. Since Wnt signaling is known to sequester these factors at the plasma membrane and to inhibit GSK3, this provides a simple and efficient mechanism to rapidly dissolve the translation factories and to stabilize β-catenin. Interestingly, one APC molecule can bind more than 30 molecules of β-catenin (Su et al., 1993). Most likely, the nascent β-catenin protein recruits APC, which can then crosslink multiple β-catenin polysomes by its ability to bind several β-catenin proteins. This suggests that the formation of the factories could follow general phase-separation mechanisms relying on multivalent interactions made by polymeric molecules: APC on one side, and β-catenin polysome on the other (which contain several nascent protein chains). Axin is also likely to contribute to the aggregation process by its ability to bind β-catenin, APC, and to oligomerize. Possibly, the β-catenin mRNAs may also directly contribute to factory formation, since they physically interact with APC (Preitner et al., 2014). While the function of APC in β-catenin metabolism is not completely understood, our data indicate a direct role in organizing β-catenin translation factories.

Overall, we found 4 translation factories while screening 500 mRNAs. Extrapolation to the 20,000 human genes suggests that few hundred such factories may exist in a cell. Translation may thus be compartmentalized to a much higher degree than previously anticipated.

## Acknowledgements

We thank Florence Rage and Susanne Schmidt for the preparation of neuronal hippocampus primary cells. We also thank the staff of MRI imaging facility for their technical support. R.C. was supported by a scholarship from the Lebanese university and the Lebanese National Council for Scientific Research (LNCSR). AdS was supported by an FRM fellowship. This project was supported by France BioImaging (ANR-10-INBS-04), the Agence Nationale de la Recherche (ANR-11-BSV8-018-02 and ANR-14-CE10-0018-01), the Institut Pasteur, the Ligue Nationale Contre le Cancer and the Fondation pour la Recherche Médicale (FRM grant to EB). This work was supported by the Labex EpiGenMed, from the framework “Investissements d’avenir”.

## Author contributions statement

EB conceived the study with help of TW, FM and KZ. Experiments were performed by RC, AdS, XP, MP, MCR, OSK, EB and AMT. AI, AuS, TW and FM designed and performed the automated image analysis. AI, AdS, XP, AH, IP, HLH, CZ, RC, FM, TW, MP, KZ, HLH and EB analyzed the data. EB prepared the Figures, with help of AdS, OSK, AI, TW, FM and RC. EB wrote the manuscript.

## STAR Methods

### Cell lines and culture conditions

The collection of HeLa-Kyoto cells stably transfected with the GFP-tagged BACs was described previously (Maliga et al., 2013; Poser et al., 2008). The BAC-GFP clones were maintained in Dulbecco’s modified Eagle’s Medium (DMEM, Gibco) supplemented with 10% fetal bovine serum (FBS, Sigma), 100 U/mL penicillin/streptomycin (Sigma) and 400 μg/ml G418 (Gibco). HeLa-Kyoto, C2C12 and NIH3T3 cell lines were grown in the same medium without G418. RPE1 Centrin2-GFP (Mikule et al., 2007; a gift of B. Delaval) were cultured in DMEM:F12 medium (Gibco) supplemented with 10% fetal bovine serum (Sigma) and 100 U/mL penicillin/streptomycin (Sigma). SH-SY5Y cells were grown in DMEM with 10% FBS and differentiated for 72h in DMEM containing 3% FBS and 10 μM retinoic acid. Drugs were used at the following final concentrations: 100 μg/ml for puromycin, 200 μg/ml for cycloheximide, 5 μg/ml for actinomycin D, 10 μM for GSK3 inhibitor CHIR99021 and 0.5 μM for LG-007. For puromycin, data are shown for 1h or treatment, but similar results were obtained after 30 minutes only (not shown). For Actinomycin D, data are shown for 1h or treatment, but similar results were obtained for up to 6h of treatment (not shown).

### Insertion of SunTag cassette by CRISPR/Cas9

The recombination cassettes contained 500 bases of homology arms flanking a puromycin resistance gene translated from the endogenous ATG sequence, followed by a P2A sequence, 32 SunTag repeats, and another P2A sequence fused to the protein of interest. Hela Kyoto cells stably expressing the scFv-GFP were transfected using JetPrime (Polyplus) and a cocktail of three plasmids, including the recombination cassette and constructs expressing Cas9-HF1 and guide

RNAs with an optimized scaffold (Chen et al., 2013). Cells were selected on 0.25 μg/ml puromycin for a few weeks. Individual clones were then picked and analyzed by PCR genotyping, fluorescent microscopy and by smiFISH with probes against the SunTag sequence. The sequences targeted by the guide RNAs were (PAM sequences are underlined): AAGTGAGCCCGACCGAGCGG<u>AGG</u> for ASPM and CGGGGTATTCGAATCGGCGG<u>CGG</u> for BUB1.

### Genotyping

PCR was done using a Platinum Taq DNA Polymerase (Invitrogen) on genomic DNA prepared with GenElute Mammalian Genomic DNA miniprep (Sigma-Aldrich). The sequences of oligonucleotides were: 5’-TGTTCCTGGAAACCGCAATG (ASPM WT forward); 5’-GTTTATGTGTTGTCCCCGCC (ASPM WT reverse); 5’-TACCCTTCTTCAGTCTGGCG (SunTag reverse).

### Treatments with siRNAs

HeLa cells were seeded on 0.17 mm glass coverslips deposited in 6-well plates. Cells were transfected at 70% confluency using JetPrime (Polyplus). Double-stranded siRNAs (30 pmoles) were diluted into 200 μl of JetPrime buffer. JetPrime reagent was added (4 μl) and the mixture was vortexed. After 10 minutes at room temperature (RT), the mix was added to the cells grown in 2 ml of serum-containing medium. After 24 hours, the transfection medium was replaced with fresh growth medium and cells were fixed 24h later. The sequences of the siRNA were: APC-1: 5’-GCACAAAGCUGUUGAAUUUdTdT-3’; APC-2: 5’-UGAAAGUGGAGGUGGGAUAdTdT-3’; APC-3: 5’-UAAUGAACACUACAGAUAGdTdT; Axin1-1: 5’-GCGUGGAGCCUCAGAAGUUdTdT; Axin1-2: 5’-CCGAGGAGAAGCUGGAGGAdTdT. The three APC siRNAs gave similar results, and the data presented correspond to the siRNA APC-1 and APC-2. Likewise, the two Axin1 gave similar results and the data presented correspond to the siRNA Axin1-1.

### Single molecule fluorescent *in situ* hybridization

Cells grown on glass coverslips were fixed for 20 min at RT with 4% paraformaldehyde diluted in PBS, and permeabilized with 70% ethanol overnight at 4°C. For smFISH, we used a set of 44 amino-modified oligonucleotide probes against the GFP-IRES-Neo sequence present in the BAC construction (sequences given in Table S3). Each oligonucleotide probe contained 4 primary amines that were conjugated to Cy3 using the Mono-Reactive Dye Pack (PA23001, GE Healthcare Life Sciences). To this end, the oligos were precipitated with ethanol and resuspended in water. For labelling, 4 μg of each probe was incubated with 6 μl of Cy3 (1/5 of a vial resuspended in 30 μl of DMSO), and 14 μl of carbonate buffer 0.1 M pH 8.8, overnight at RT and in the dark, after extensive vortexing. The next day, 10 μg of yeast tRNAs were added and the probes were precipitated several times with ethanol until the supernatant lost its pink color. For hybridization, fixed cells were washed with PBS and hybridization buffer (15% formamide in 1xSSC), and then incubated overnight at 37°C in the hybridization buffer also containing 130 ng of the probe set for 100 μl of final volume, 0.34 mg/ml tRNA (Sigma), 2 mM VRC (Sigma), 0.2 mg/ml RNAse-free BSA (Roche Diagnostic), and 10% Dextran sulfate. The next day, the samples were washed twice for 30 minutes in the hybridization buffer at 37°C, and rinsed in PBS. Coverslips were then mounted using Vectashield containing DAPI (Vector laboratories, Inc.).

For smiFISH, 24 to 48 unlabeled primary probes were used (sequences given in Table S3). In addition to hybridizing to their targets, these probes contained a FLAP sequence that was hybridized to a secondary fluorescent oligonucleotide. To this end, 40 pmoles of primary probes were pre-hybridized to 50 pmoles of secondary probe in 10 μl of 100 mM NaCl, 50 mM Tris-HCl, 10 mM MgCl^2^, pH 7.9. Hybridization was performed at 85°C for 3 min, 65°C for 3 min, and 25°C for 5 min. The final hybridization mixture contained the probe duplexes (2 μl per 100 μl of final volume), with 1X SSC, 0.34 mg/ml tRNA (Sigma), 15% Formamide, 2 mM VRC (Sigma), 0.2 mg/ml RNAse-free BSA, 10% Dextran sulfate. Slides were then processed as above. For AKAP9, the probes used were RNA and not DNA (sequence in Table S3). The protocol was similar except that hybridization was performed at 48°C and that 50 ng of primary probe (total amount of the pool of probes) and 30 ng of the secondary probes were used per 100 μl of hybridization mix.

### Immunofluorescence

HeLa cells were seeded and fixed as for smFISH. Cells were permeabilized with 0.1% Triton-X100 in PBS for 10 minutes at RT and washed twice with PBS. For P-body labelling, coverslips were incubated overnight at 4°C with a monoclonal mouse antibody (Santa Cruz sc-8418), diluted 1/250 in PBS containing 0.1% BSA. Coverslips were washed three times with PBS, 10 minutes each time, and incubated with an FITC anti-mouse secondary antibody (Jackson ImmunoResearch 115096-006) diluted 1/200 in PBS with 0.1% BSA. After 1 hour of incubation at RT, coverslips were washed three times with PBS, 10 minutes each. For APC labelling, coverslips were incubated 1h at RT with an anti-APC mouse monoclonal antibody (F3, Santa Cruz sc-9998), diluted 1/50 in PBS containing 0.1% BSA. For the other epitopes, the antibodies were: (i) N-terminus of β-catenin (Cell Signaling #9581 diluted 1/40, or Cell Signaling #2698 diluted 1/400); (ii) S33/S37 phospho β-catenin (Cell Signaling #9561 diluted 1/800); (iii) conjugated ubiquitin (Viva Bioscience mAB FK2 diluted 1/100); (iv) Axin1 (Cell Signaling #2087 at 1mg/ml, diluted 1/200). After washing as above, coverslips were incubated 1h at RT with a secondary antibody diluted in PBS with 0.1% BSA. Coverslips were washed again and mounted using Vectashield containing DAPI (Vector laboratories, Inc.). The secondary antibodies were as follow: goat anti-mouse FITC (Jackson 115-095-062 diluted 1/200); goat anti-rabbit FITC (Jackson 111-096-047 diluted 1/200); donkey antirabbit Cy3 Jackson (711-165-152 diluted 1/200); goat anti-mouse Cy3 Jackson (115-165-003 diluted 1/200); goat anti-mouse Cy5 (Jackson 115-176-003 diluted 1/400); goat anti-rabbit Cy5 (Jackson 111-175-144 diluted 1/400).

To label the translation machinery, HeLa cells were rinsed with PBS and fixed by 4 % paraformaldehyde (PFA) for 10 minutes in room temperature (RT). After fixation, cells were washed with PBS and permeabilized through 0.1 % Triton X-100 (EUROMEDEX, 2000-A) for 3 minutes in RT. After washing with PBS, cells were incubated with 1 % Bovine Serum Albumin (BSA, Sigma Aldrich, A-7906) in PBS for 30 minutes in RT. Primary antibodies against FOP (Abnova, H00011116-M01), eIF4E (Santa Cruz, SC-13963), phospho-RPS6 (Santa Cruz, SC-54279) were incubated with cells for 1 hour in RT. HOECHST 33258 (Sigma-Aldrich) was incubated for 5 minutes in RT and secondary antibodies fused with Alexa 488 (Thermofisher, A21141) and Cy3 (Jackson Immuno Research, 111-165-144) were incubated for 1 hour in RT. Coverslisps were mounted with Fluoromount-G^tm^ (Invitrogen 00-4958-02).

### Imaging of fixed cells

Microscopy slides were imaged on: (i) a Zeiss AxioimagerZ1 wide-field microscope equipped with a motorized stage, a camera scMOS ZYLA 4.2 MP, using a 63x or 100x objective (Plan Apochromat; 1.4 NA; oil); or (ii) a Nikon Ti fluorescence microscope equipped with ORCA-Flash 4.0 digital camera (HAMAMATSU), for the eIF4E and p-RPS6 IFs. Images were taken as z-stacks with one plane every 0.3 μm. The microscope was controlled by MetaMorph and Figures were constructed using ImageJ, Adobe Photoshop and Illustrator.

### Imaging of live cells

Live imaging was done using a spinning disk confocal microsope (Nikon Ti with a Yokogawa CSU-X1 head) operated by the Andor iQ3 software. Acquisitions were performed using a 100X objective (Nikon CF1 PlanApo λ 1.45 NA oil), and an EMCCD iXon897 camera (Andor). Samples were sequentially excited at 488 and 640nm. Cells were maintained in anti-bleaching live cell visualization medium (DMEM^gfp^; Evrogen), supplemented with 10% fetal bovine serum at 37°C in 5% CO2. SiR-DNA (Spirochrome) was kept at 100 nM throughout the experiments to label DNA.

### Deconvolution

For Figure S8A and S8B, deconvolution was performed using the Huygens Professional software (Scientific Volume Imaging, Hilversum, Netherlands). The point spread function (PSF) was a theoritical one, and background values were manually estimated. The deconvolution was done using the CMLE deconvolution algorithm, with 50 iterations and a quality threshold of 0.01, and without bleaching correction.

### Image analysis and quantifications

Nuclear segmentation was performed from the DAPI channel, cell segmentation from the autofluorescence of the actual smFISH image. Prior to segmentation, 3D images were projected to 2D images as previously described (Tsanov et al., 2016). Nuclei were segmented with a Deep Neural Network (Matterport implementation of the Mask R-CNN, He et al., 2017), trained on the Data Science Bowl 2018. Cells were segmented with a custom algorithm, based on prefiltering the 2D projected smiFISH image, followed by a watershed based method using nuclear regions as seeds. In case of poor results, segmentation was corrected manually. RNAs were detected with a Python implementation of FISH-quant (Mueller et al., 2013), by applying a local maximum detection on Laplacian of Gaussian (LoG) filtered images, and by decomposing larger agglomerations of spots as previously described (Samacoits et al., 2018). The quantification of the number of mRNA per cell is shown in Figure S5D.

To analyze the percentage of mRNAs molecules in protrusions, we first detected protrusions by calculating the difference between the segmented cellular region and its opening with a size of 30. The percentage of RNAs detected in the residuum was then divided by the total number of cytoplasmic mRNAs. Enrichment values were calculated with respect to a uniform distribution of mRNAs molecules.

Foci were detected with the DBSCAN algorithm on the detected spots (Ester et al., 1996). We defined foci as sets of at least 5 points where for each point there is at least one other point closer than 350 nm that is also part of the focus. Percentages of RNA in foci were then calculated as number of RNA inside the foci divided by the number of cytoplasmic RNAs.

To measure the degree of spatial overlap between mRNA (by smFISH) and protein (by the GFP fluorescence), we calculated an enrichment ratio. Cells and nuclei were outlined manually in 2D based on the GFP and DAPI image, respectively. The subsequent analysis was restricted to the cytoplasm. FISH-quant was used to detect mRNAs in 3D and each cell was post-processed separately. First, we calculated the median pixel intensity in the GFP image at the identified RNA positions. Second, we estimated a normalization factor as the median GFP intensity of the outlined cytoplasm within the z-range of the detected mRNAs. The enrichment ratio was then estimated as the ratio of the median GFP intensity at the RNA positions divided by the mean cytoplasmic GFP intensity.

### Automated classification of RNA localization patterns

From the segmentation results and the spot locations, we filtered out cells with less than 30 mRNAs and then calculated the following features, which describe the spatial distribution of points inside the cell: (i) average mRNA distance to the cytoplasmic membrane (normalized to the value obtained under a uniform distribution); (ii) average mRNA distance to the nuclear membrane (normalized to the value obtained under a uniform distribution); (iii) proportion of RNA inside the nucleus; (iv) number of foci (detected as described above); (v) proportion of RNA inside foci; (vi) average foci distance to the cytoplasmic membrane (normalized to the value obtained under a uniform foci distribution); (vii) average foci distance to the nuclear membrane (normalized to the value obtained under a uniform foci distribution); (viii) peripheral dispersion index, defined as the expectation of the squared point distance to the centroid of the cell; (ix) proportion of RNA inside protrusion (detected as described above); (x) number of mRNA within 515 nm from the nuclear membrane (normalized to the value obtained under a uniform distribution); (xi) number of RNA between 515 nm and 1030 nm from the nuclear membrane (normalized to the value obtained under a uniform distribution); (xii) number of RNA between 1030 nm and 1545 nm from the nuclear membrane (normalized to the value obtained under a uniform distribution); (xiii) number of RNA between 0 and 515 nm from the cytoplasmic membrane (normalized to the value obtained under a uniform distribution); (xiv) number of RNA between 515 nm and 1030 nm from the cytoplasmic membrane (normalized to the value obtained under a uniform distribution); (xv) number of RNA between 1030 nm and 1545 nm from the cytoplasmic membrane (normalized to the value obtained under a uniform distribution).

In order to visually explore the structure in the data, we perform t-Distributed Stochastic Neighbor Embedding (t-SNE; Van der Maaten et al., 2008). Perplexity was chosen to 30 according to their guidelines.

For manual annotation, we generated small panels with the original single cell images and the segmentation and RNA detection results. These images were then classified manually into different class corresponding to their localization patterns. The manual classification result was then independently checked by several trained microscopists. The manual annotations were visualized in the t-SNE plot and used as a training set for training an automated classifier for the recognition of RNA localization patterns.

For supervised learning, we trained 5 binary Random Forest classifiers (Breiman L., 1999), where the actual training set (320 to 750 cells) was composed of all the cells of one class and a subsampling of cells from the union of all other classes such that the imbalance was 1:5. This strategy is known as “one vs. all”. Importantly, the classifiers were independent, and for this reason allowed to assign several patterns to the same cell. Training parameters were 100 trees with a maximum depth of 3 and a minimum number of samples per splitter node of 2. For each split, we consider a subset of 10 features and entropy criterion. Percentages were then calculated and visualized in a heatmap. P-values were calculated with Fisher’s exact test. For the single-gene heatmap visualizations, we assigned to each cell a vector of classification probabilities. Note that the vectors do not sum to one because classifiers are independent. The classification vectors were then hierarchically clustered using Average method with Euclidean distance. All visualizations were done with Matplotlib. The code was written in Python and together with the preprocessed image data is accessible at GitHub (https://github.com/Henley13/paper_translation_factories_2020). For t-SNE and Random Forests we used the scikit-learn implementations (Pedregosa et al., 2011), and for basic image analysis scikit-image (Van der Walt et al., 2014). All the pattern quantifications were done using the random forest classifiers except for figures 6B, 7D, and 9E-F, where cells were manually annotated to assign them a pattern type.

## Supplemental Spreadsheets and Movie

**Table S1, related to Figure 2 and 4.** Summary of all the mRNAs screened.

**Table S2, related to Figure 2 and 4.** Summary of all the localized mRNAs.

**Table S3, related to all Figures.** Sequence of the smFISH and smiFISH probes.

**Movie, related to Figure 6:** SunTagged ASPM polysomes (green) are anchored on the nuclear envelope. DNA (red) is stained with SiR-DNA. The movie is made from a maximal image projection of z-stacks, acquired every 40 seconds for 50 minutes.

