## Supplementary Information for "A localization screen reveals translation factories and widespread co-translational RNA targeting"

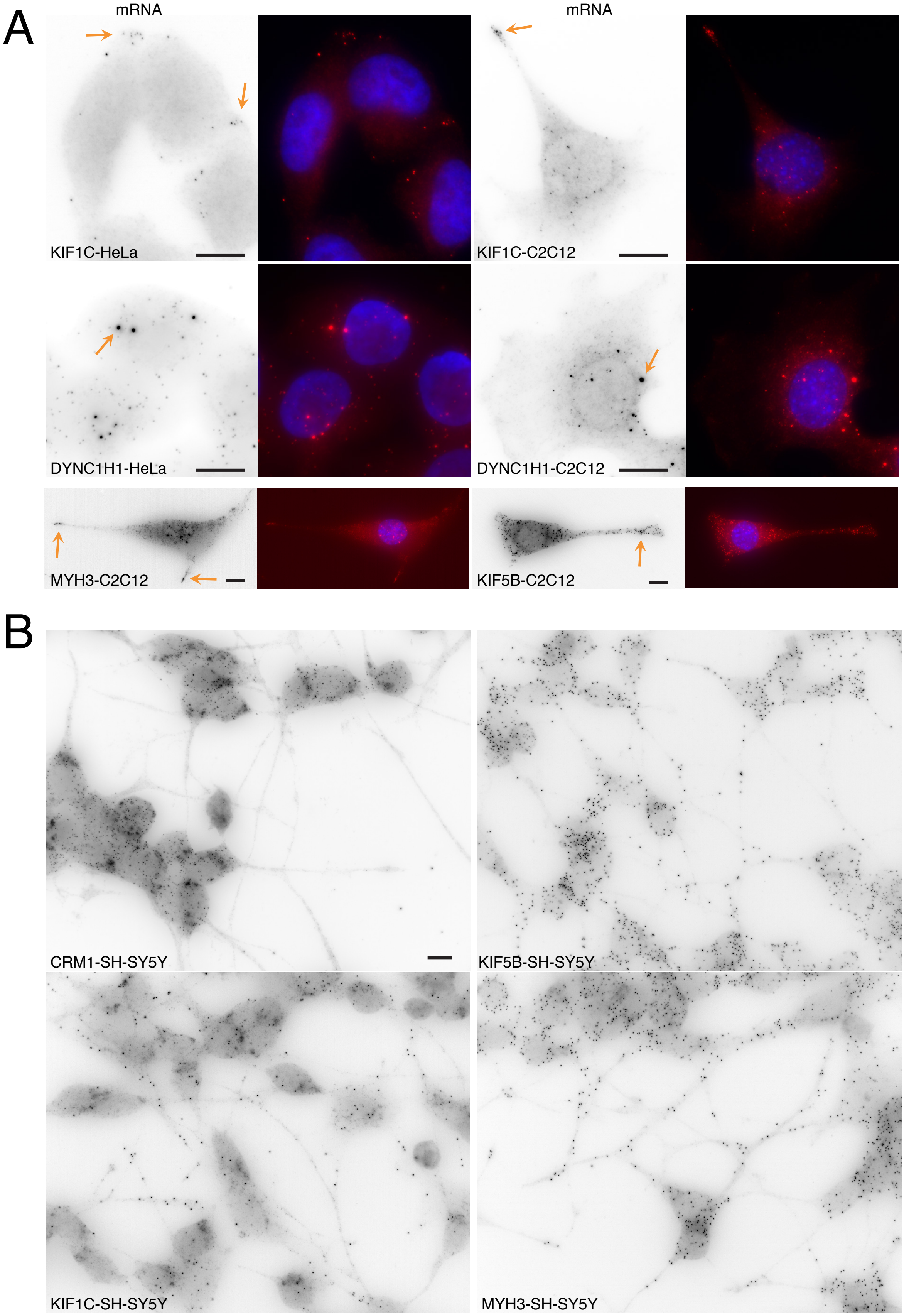
A
mRNA
KIF1C-HeLa
mRNA
KIF1C-C2C12
DYNC1H1-HeLa
DYNC1H1-C2C12
MYH3-C2C12
KIF5B-C2C12
B
CRM1-SH-SY5Y
KIF5B-SH-SY5Y
KIF1C-SH-SY5Y
MYH3-SH-SY5Y

Figure S1

**Figure S1, related to Figure 2. Several endogenous mRNAs coding for motor proteins localize in diverse cell lines.**

A-KIF1C, DYNC1H1, MYH3 and KIF5B mRNAs in HeLa and mouse C2C12 cells. Images are micrographs of the indicated cell line, hybridized with Cy3-labelled oligonucleotide probes against the indicated endogenous mRNAs. Red and black: signal from the probes. Blue: DAPI staining. Arrows point to mRNA localizing in foci (DYNC1H1), or at the cell periphery. Scale bars: 10 microns.

B-CRM1, KIF1C, KIF5B and MYH3 mRNAs in differentiated SH-SY5Y neuroblast cells. Legend as in A; scale bar: 10 microns.

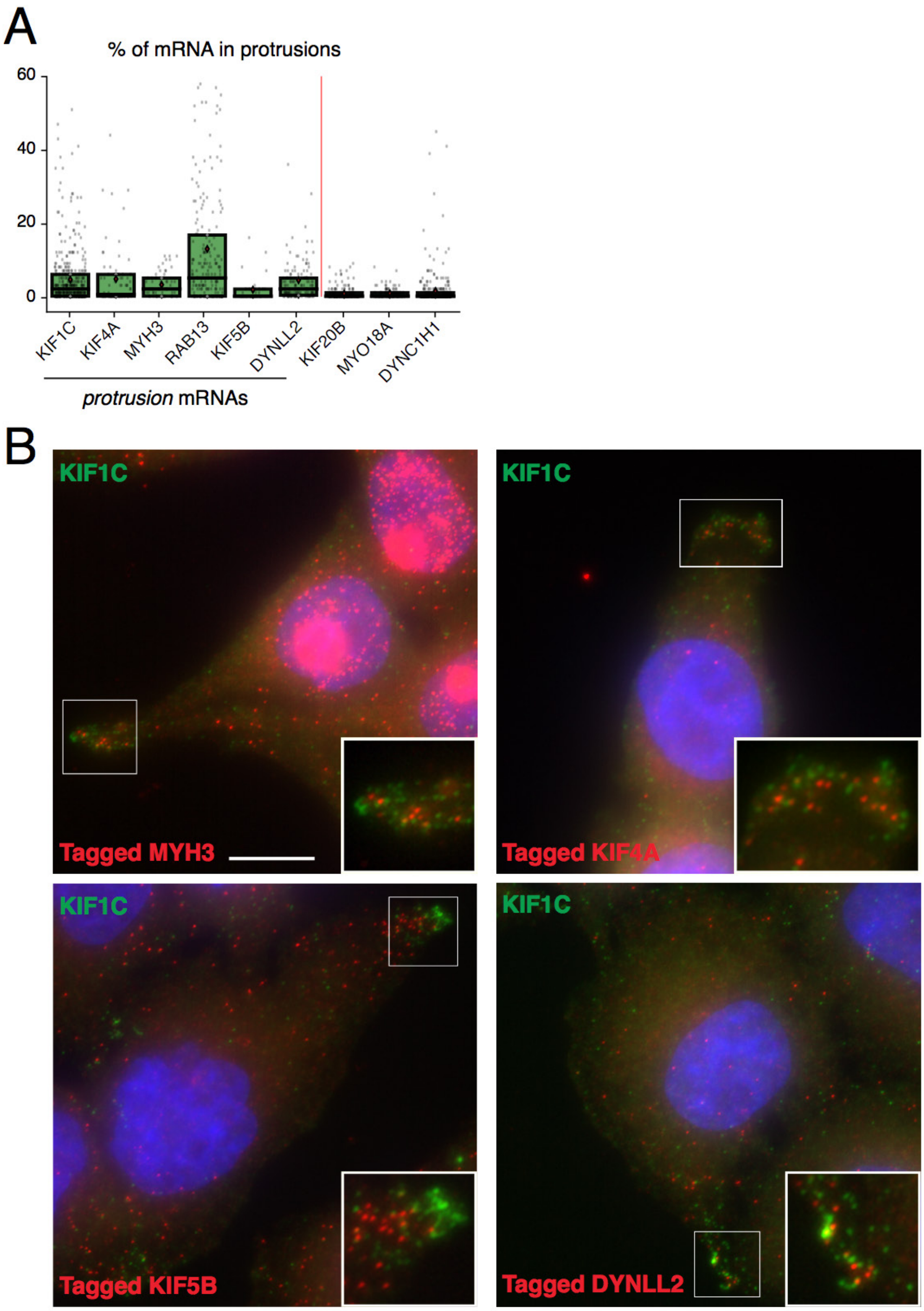
A
% of mRNA in protrusions
60
40
20
0
KIF1C
KIF4A
MYH3
RAB13
KIF5B
DYNLL2
KIF20B
MYO18A
DYNC1H1
*protrusion* mRNAs
B
KIF1C
Tagged MYH3
KIF1C
Tagged KIF4A
KIF1C
Tagged KIF5B
KIF1C
Tagged DYNLL2

Figure S2

**Figure S2, related to Figure 2 and Figure 3. Localization of mRNAs in cytoplasmic protrusions.**

A-Quantification of the amount mRNA accumulating in cytoplasmic protrusions. Left: the bar plot depicts the fraction of mRNA molecules localizing in cytoplasmic protrusion. Each cell is a spot, and the box corresponds to the second and third quartiles; the black bar is the median and the red diamond the mean.

B-Co-localization of mRNAs in cytoplasmic protrusions. Images are micrographs of HeLa cells hybridized with Cy3- and Cy5-labelled oligonucleotide probes against the indicated mRNAs (red and green, respectively). Blue: DAPI staining. Scale bar: 10 microns. Insets: zoom of the boxed areas.

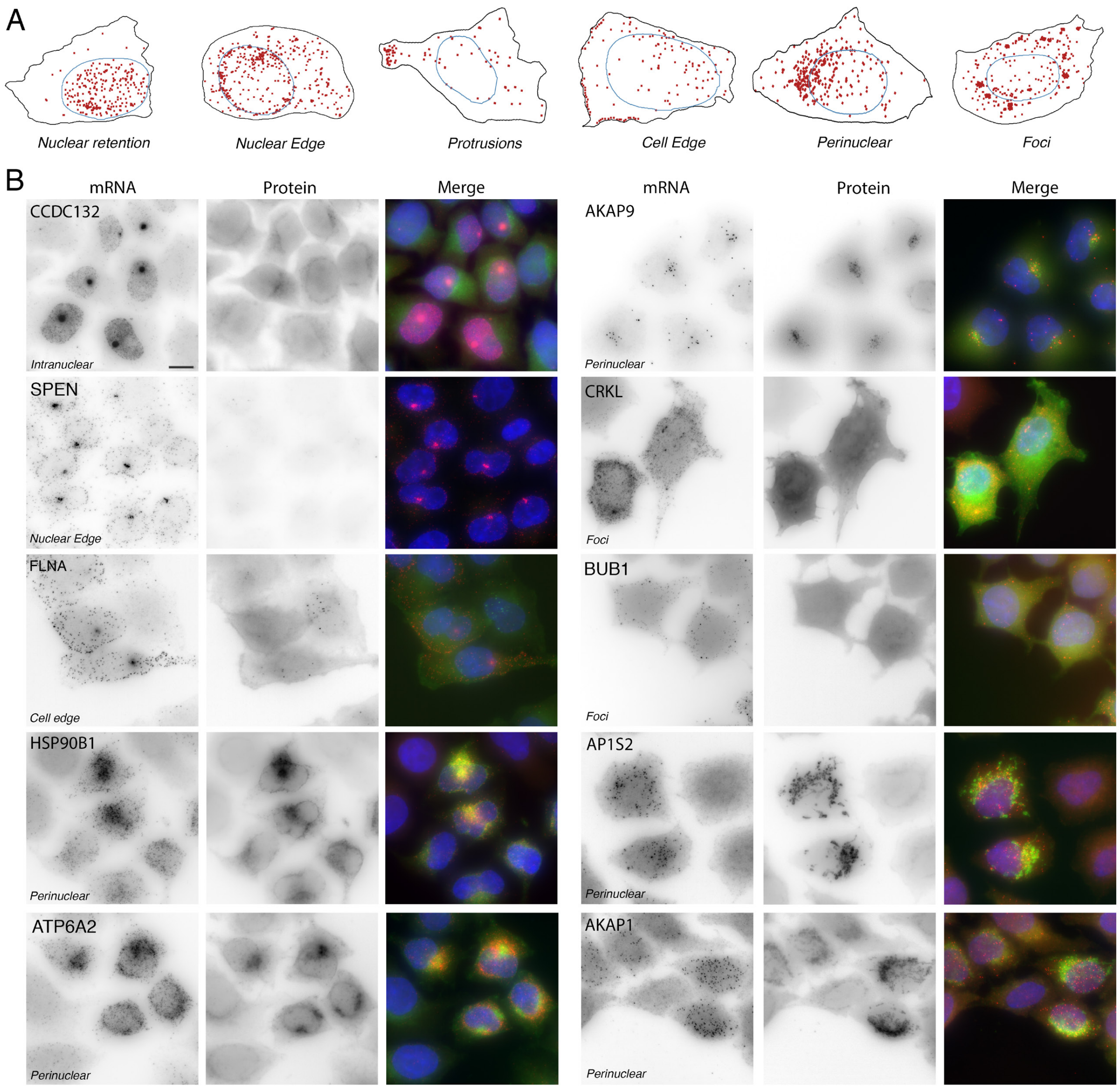


**Figure S3, related to Figure 3. Examples of localized mRNAs found in the BAC screen.**

A-Schematic of the localization classes. The mRNAs are in red; nuclei are delimited by blue lines and cellular areas by black lines.

B-Localized tagged mRNAs. Legend as in Figure 1B. The tagged gene on the BAC is indicated (top of each mRNA panel), as well as the localization class (italics). Scale bar: 10 microns.

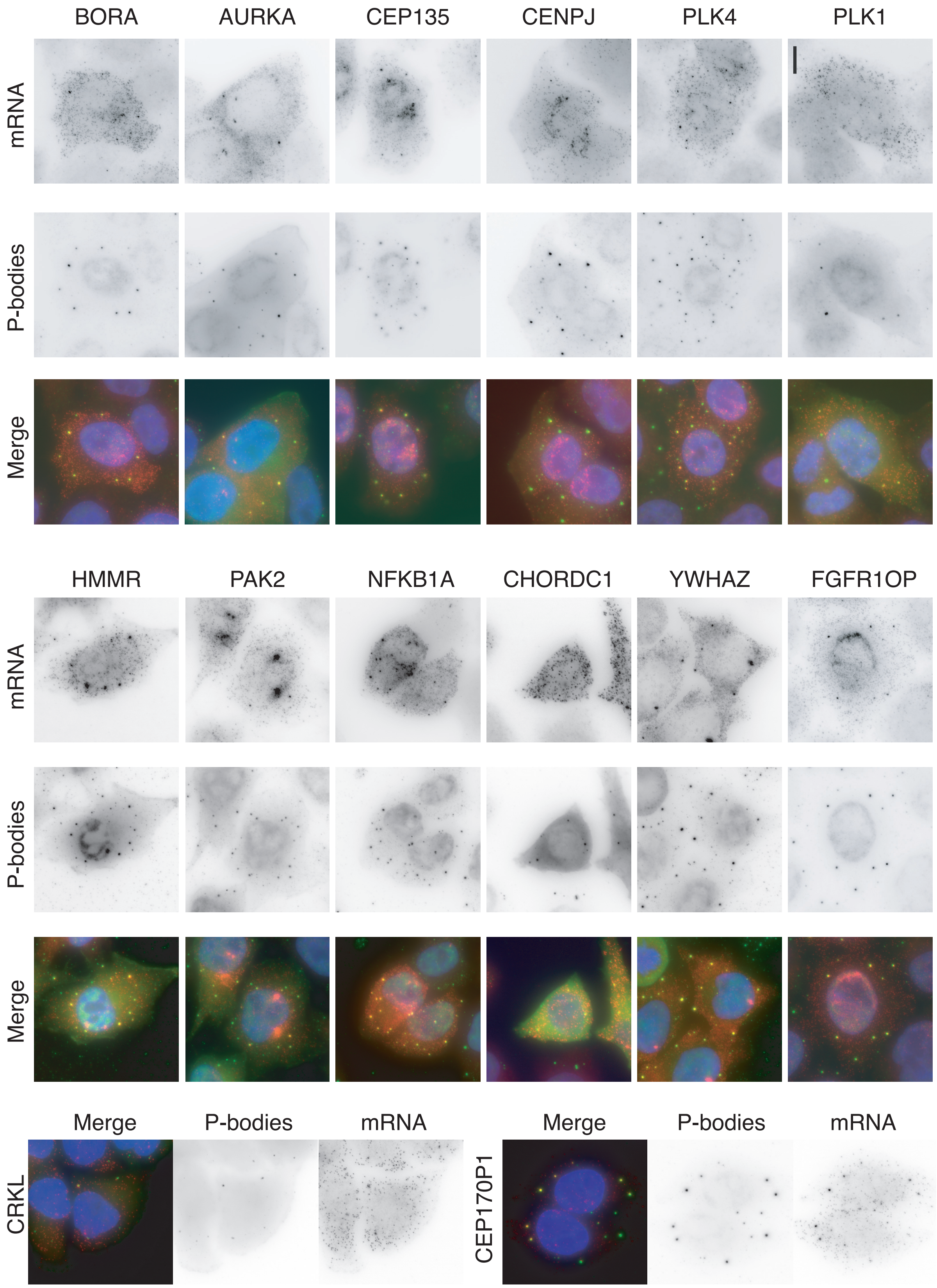
BORA
AURKA
CEP135
CENPJ
PLK4
PLK1
mRNA
P-bodies
Merge
HMMR
PAK2
NFKB1A
CHORDC1
YWHAZ
FGFR1OP
mRNA
P-bodies
Merge
Merge
P-bodies
mRNA
CRKL
Merge
P-bodies
mRNA
CEP170P1

Figure S4

**Figure S4, related to Figure 3 and 4. BAC-tagged mRNAs accumulating in P-bodies.** Images are micrographs of HeLa cells expressing the indicated tagged BAC gene, and both hybridized with Cy3-labelled probes against the tag (left and green in the merge), and labelled by indirect immuno-fluorescence with an antibody against P-bodies (middle panel and green in the merge). Scale bar: 10 microns.

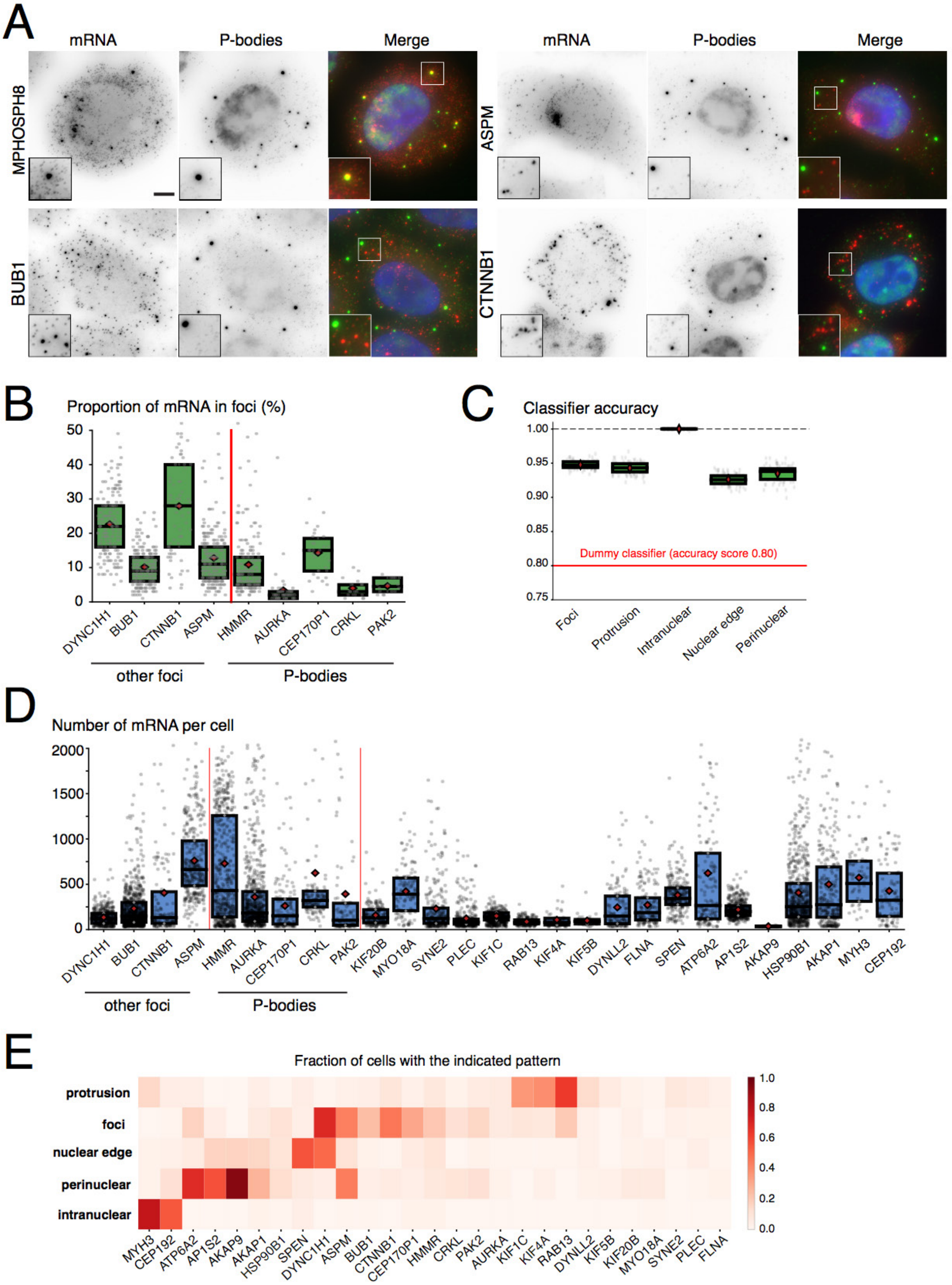

A
mRNA
P-bodies
Merge
MPHOSPH8
ASPM
BUB1
CTNNB1
B
Proportion of mRNA in foci (%)
DYNC1H1
BUB1
CTNNB1
ASPM
HMMR
AURKA
CEP170P1
CRKL
PAK2
other foci
P-bodies
C
Classifier accuracy
Dummy classifier (accuracy score 0.80)
Foci
Protrusion
Intranuclear
Nuclear edge
Perinuclear
D
Number of mRNA per cell
other foci
P-bodies
E
Fraction of cells with the indicated pattern
protrusion
foci
nuclear edge
perinuclear
intranuclear


Figure S5

**Figure S5, related to Figure 3 and 5. Quantitative analysis of mRNA localization.**

A-Foci containing BUB1, ASPM, or CTNNB1 mRNAs do not co-localize with P-bodies. The experiment was performed with the BAC-tagged cell lines. Legend as in A. Scale bar: 10 microns.

B-Fraction of mRNA in foci. Each cell is a spot, and the box correspond to the 2$^{nd}$ and 3$^{d}$ quartiles, the bar is the median and the red diamond is the mean. Only cells classified as *'foci'* by the Random Forest classifier are considered.

C-Classifier accuracy. The values depicts the fraction of correct classifier predictions to the total number of input values. The dummy classifier always predicts "no pattern".

D- Total number of mRNA per cell. Legend as in B.

E- Heatmap depicting the fraction of cells classified in the indicated pattern, for the different genes analyzed by the automated pipeline.

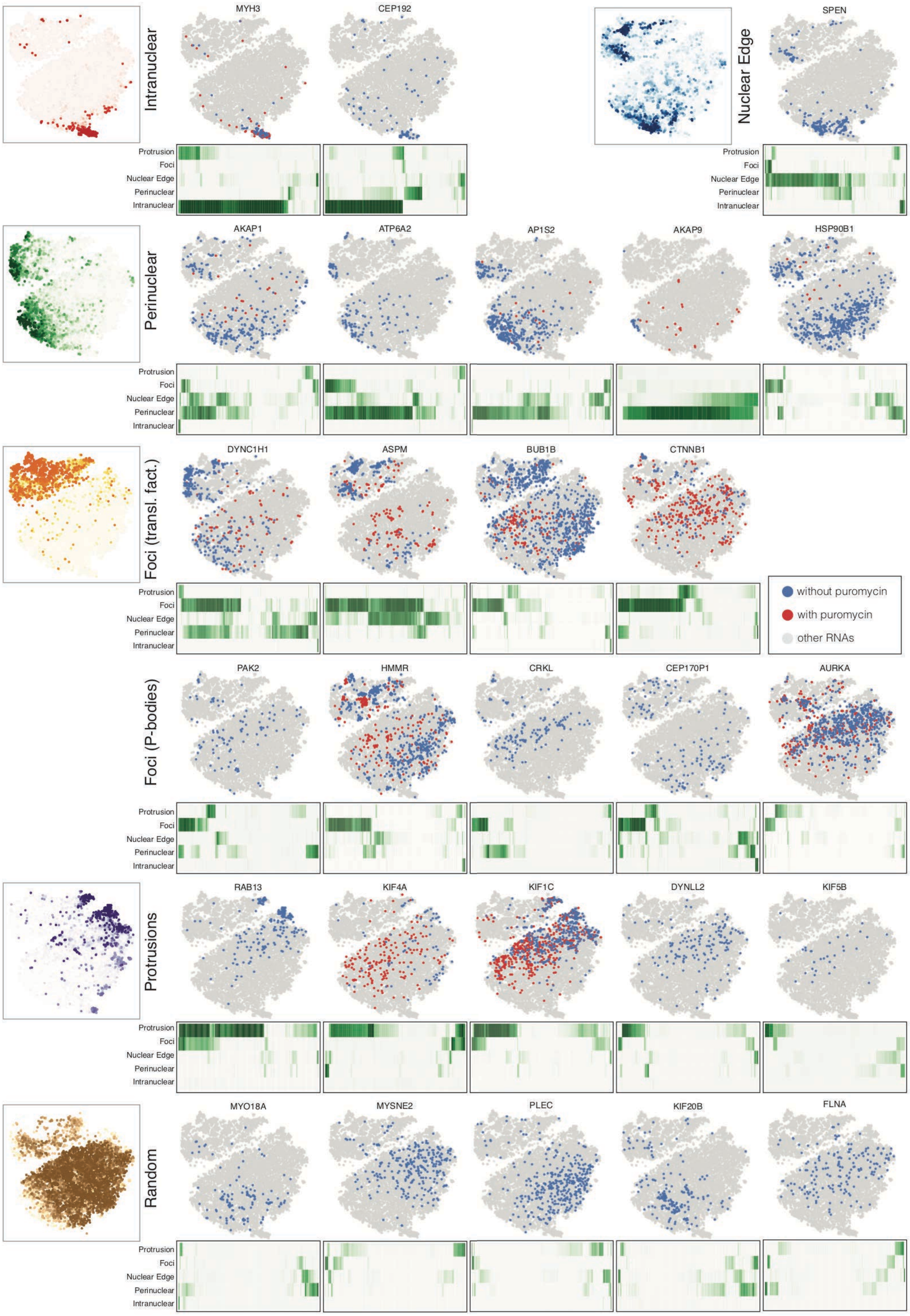

Intranuclear
MYH3
CEP192
Nuclear Edge
SPEN
Perinuclear
AKAP1
ATP6A2
AP1S2
AKAP9
HSP90B1
Foci (transl. fact.)
DYNC1H1
ASPM
BUB1B
CTNNB1
without puromycin
with puromycin
other RNAs
Foci (P-bodies)
PAK2
HMMR
CRKL
CEP170P1
AURKA
Protrusions
RAB13
KIF4A
KIF1C
DYNLL2
KIF5B
Random
MYO18A
MYSNE2
PLEC
KIF20B
FLNA
Protrusion
Foci
Nuclear Edge
Perinuclear
Intranuclear


Figure S6

**Figure S6, related to Figure 4 and 5. Quantitative analysis of mRNA localization.**

Genes are arranged in rows according to their localization class, and for each gene a t-SNE plot visualizes each cell of that gene in the embedded feature space (top), with untreated cells in blue and puromycin-treated cells in red. Below, a heatmap depicts the Random Forest classification probability of the untreated cells for each of the localization pattern (bottom). Left, a t-SNE plot with cells colored according to the probability assigned by the classifier corresponding to the gene class; color code as in 5B.

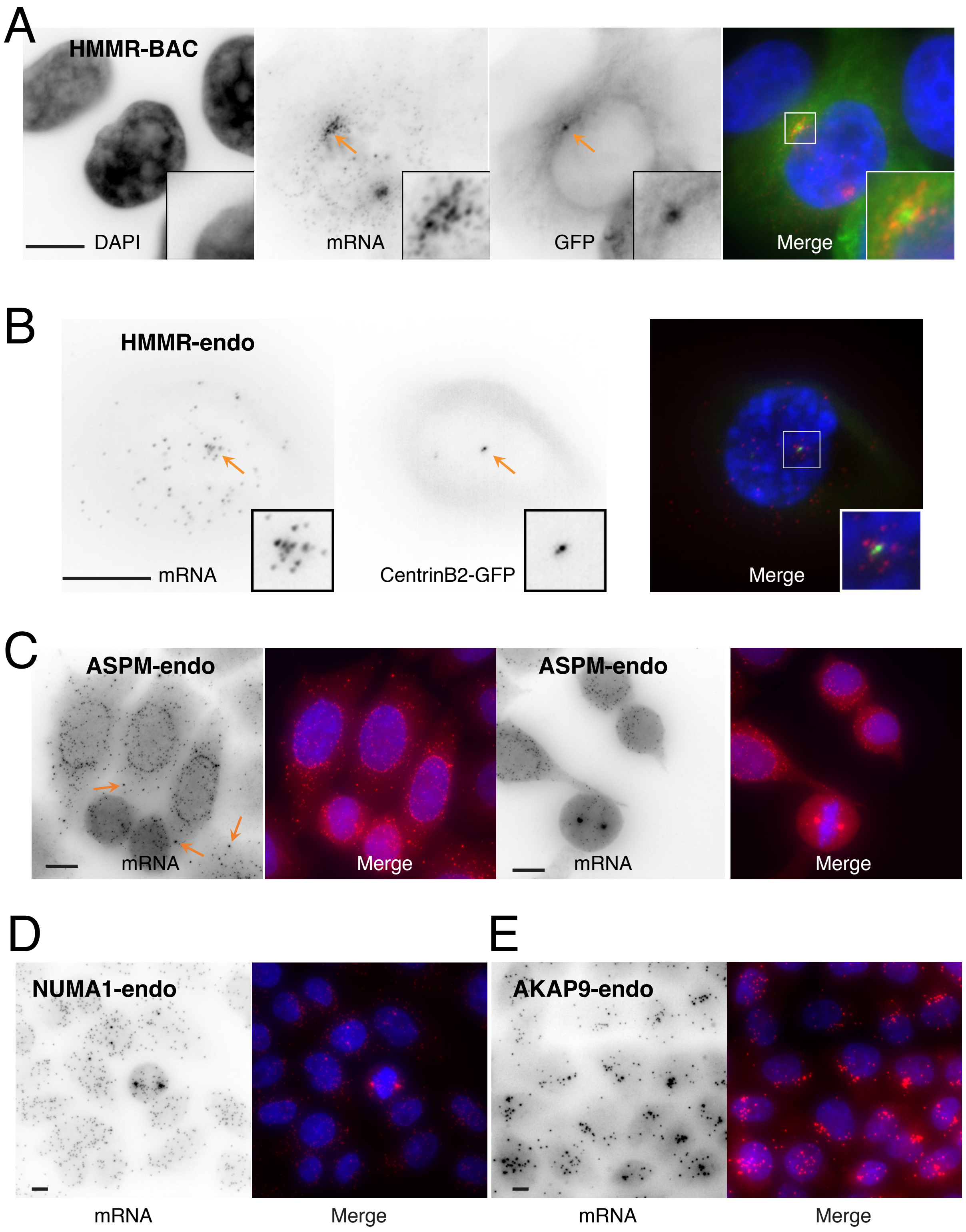

A
HMMR-BAC
DAPI
mRNA
GFP
Merge
B
HMMR-endo
mRNA
CentrinB2-GFP
Merge
C
ASPM-endo
mRNA
Merge
ASPM-endo
mRNA
Merge
D
NUMA1-endo
E
AKAP9-endo
mRNA
Merge
mRNA
Merge


Figure S7

**Figure S7, related to Figure 4. Localization of endogenous mRNAs at the centrosome.**

A-Tagged HMMR mRNA accumulates in the peri-centrosomal region. Legend as in Figure 1B. Arrows point the centrosome. Scale bar: 10 microns.

B-Endogenous HMMR mRNAs accumulate at the centrosome. Images display micrographs of RPE1 cells stably expressing Centrin2-GFP (middle and green in the merge), and hybridized with Cy3-labelled probes hybridizing to the endogenous HMMR mRNA. Scale bar: 10 microns. Arrows point the centrosome; inset: zoom of the boxed area.

C-Endogenous ASPM mRNAs accumulate at the nuclear envelope and at the spindle poles. Images display micrographs of HeLa cells hybridized with Cy3-labelled oligonucleotides against endogenous ASPM mRNAs (black and red in the merge). Orange arrows: mRNA foci. Scale bar: 10 microns.

D-Endogenous NUMA1 mRNAs accumulate at the spindle poles. Legend as in C except that probes hybridized to NUMA1 mRNAs.

E-Endogenous AKAP9 mRNAs cluster around the nucleus. Legend as in C except that probes hybridized to AKAP9 mRNAs.

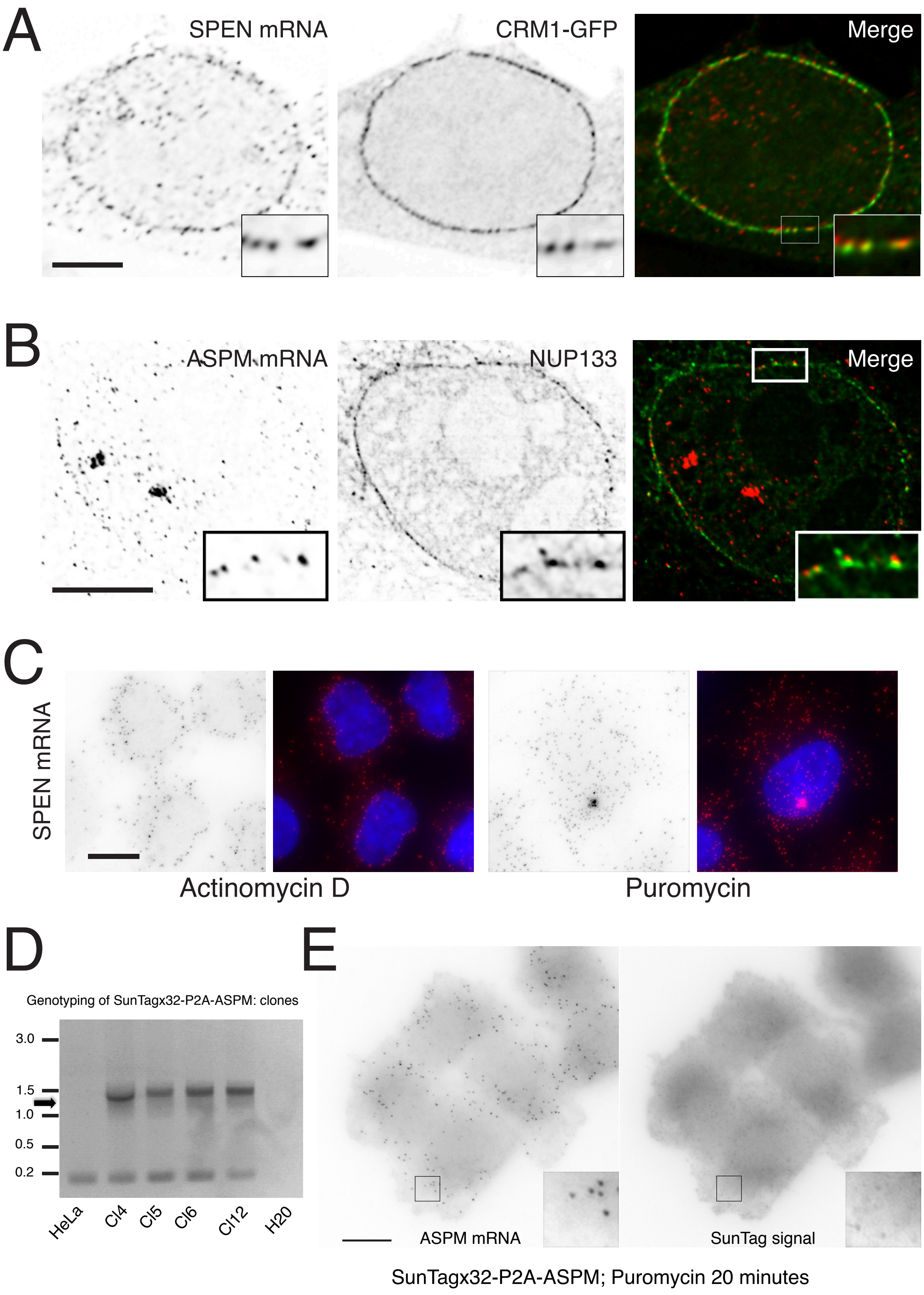
A
SPEN mRNA
CRM1-GFP
Merge
B
ASPM mRNA
NUP133
Merge
C
SPEN mRNA
Actinomycin D
Puromycin
D
Genotyping of SunTagx32-P2A-ASPM: clones
3.0
1.5
1.0
0.5
0.2
HeLa
Cl4
Cl5
Cl6
Cl12
H20
E
ASPM mRNA
SunTag signal
SunTagx32-P2A-ASPM; Puromycin 20 minutes

Figure S8

**Figure S8, related to Figure 6. Localization of BAC-tagged SPEN and ASPM mRNAs at nuclear envelope.**

A-B-Labelling of BAC-tagged SPEN (A) and ASPM (B) mRNAs with the nuclear pores. Panels are deconvolved images of HeLa cells expressing the indicated tagged BAC gene and both hybridized with Cy3-labelled oligonucleotides against the GFP-IRES-Neo tag (left and red in the merge), and labelled for nuclear pores with a transfected CRM1-GFP construct (A; middle and green in the merge) or with an antibody against NUP133 (B; middle and red in the merge). Scale bar: 10 microns.

C-Puromycin, but not Actinomycin D, disrupts localization of tagged SPEN mRNAs at the nuclear envelope. Legend as in S7A, with cells treated for 30 minutes with the indicated drug. In the puromycin experiment, the RNA foci is a transcription site. Scale bar: 10 microns.

D-Genotyping of SunTagged ASPM clones. The image shows a gel loaded with PCR reactions on genomic DNA of SunTagged HeLa clones, with primers detecting the recombined allele. HeLa: parental HeLa DNA; H2O: control without template. The arrow points the expected size (1.3 kb), and a molecular weight marker is shown on the left. Clone 4 was used in subsequent experiments.

E-Puromycin disrupts the ASPM SunTag spots colocalizing with ASPM mRNAs. Images are micrographs of HeLa cells carrying a SunTagged ASPM allele, and hybridized with Cy3-labelled probes against the SunTag sequence (left). Insets: zoom of the boxed area. Scale bar: 10 microns.

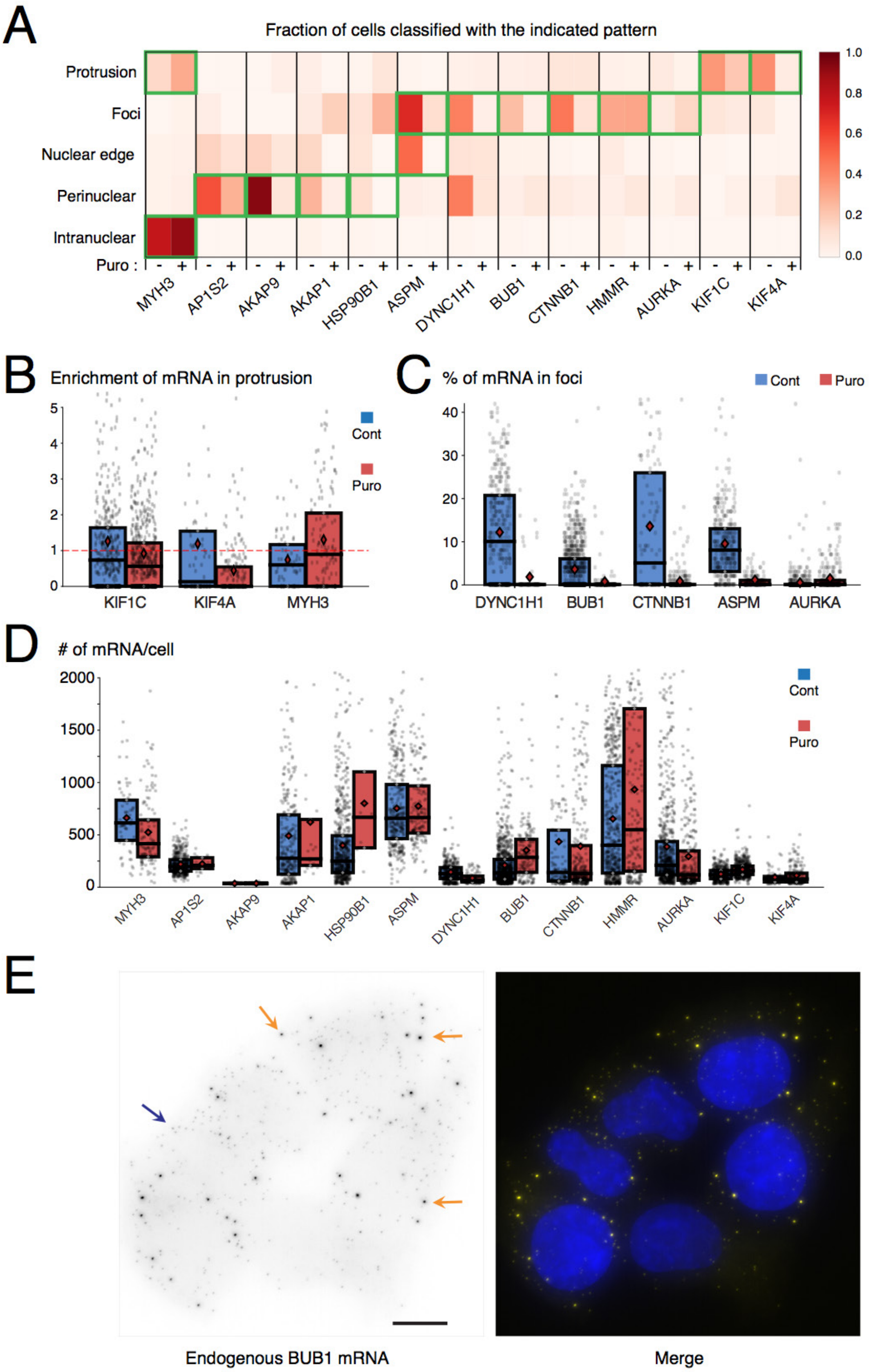
A
Fraction of cells classified with the indicated pattern
Protrusion
Foci
Nuclear edge
Perinuclear
Intranuclear
Puro :
MYH3
AP1S2
AKAP9
AKAP1
HSP90B1
ASPM
DYNC1H1
BUB1
CTNNB1
HMMR
AURKA
KIF1C
KIF4A
B
Enrichment of mRNA in protrusion
Cont
Puro
KIF1C
KIF4A
MYH3
C
% of mRNA in foci
Cont
Puro
DYNC1H1
BUB1
CTNNB1
ASPM
AURKA
D
### of mRNA/cell
Cont
Puro
E
Endogenous BUB1 mRNA
Merge

Figure S9

**Figure S9, related to Figure 8. Puromycin disrupts mRNA localization**

A-Automated classification of mRNA localization after puromycin treatment. The graph is a heatmap depicting the fraction of cells classified in the indicated pattern, for the different genes analyzed by the automated pipeline, with and without puromycin treatment for 1h. The cases boxed in green correspond to the manual annotations. For HMMR, note that in presence of puromycin the mRNA still localized to P-bodies although it is no longer at centrosomes, explaining why the automated classifier still count a similar number of cells with the pattern *'foci'*.

B-Enrichment of mRNA in protrusions after puromycin treatment (1h). This value is calculated by dividing the fraction of mRNA in protrusion by the fraction of the surface occupied by the protrusion. Each cell is a spot, and the box corresponds to the $2^{nd}$ and $3^{d}$ quartiles, the bar is the median and the red diamond is the mean.

C-Fraction of mRNAs in foci after puromycin treatment (1h). This value is calculated by dividing the number of molecules in foci by the total number of mRNA molecules in the cytoplasm. Each cell is a spot, and the box corresponds to the $2^{nd}$ and $3^{d}$ quartiles, the bar is the median and the red diamond is the mean.

D-Total number of mRNA per cell in presence and absence of puromycin treatment. Each cell is a spot, and the box correspond to the $2^{nd}$ and $3^{d}$ quartiles, the bar is the median and the red diamond is the mean. Blue: no treatment; red: puromycin for 1h.

E-Endogenous BUB1 mRNA accumulates in foci. Images are micrographs of HeLa cells hybridized with Cy3-labelled probes against BUB1 mRNA (black and yellow in the merge). Orange arrows: mRNA foci; blue arrow: single mRNA. Scale bar: 10 microns.

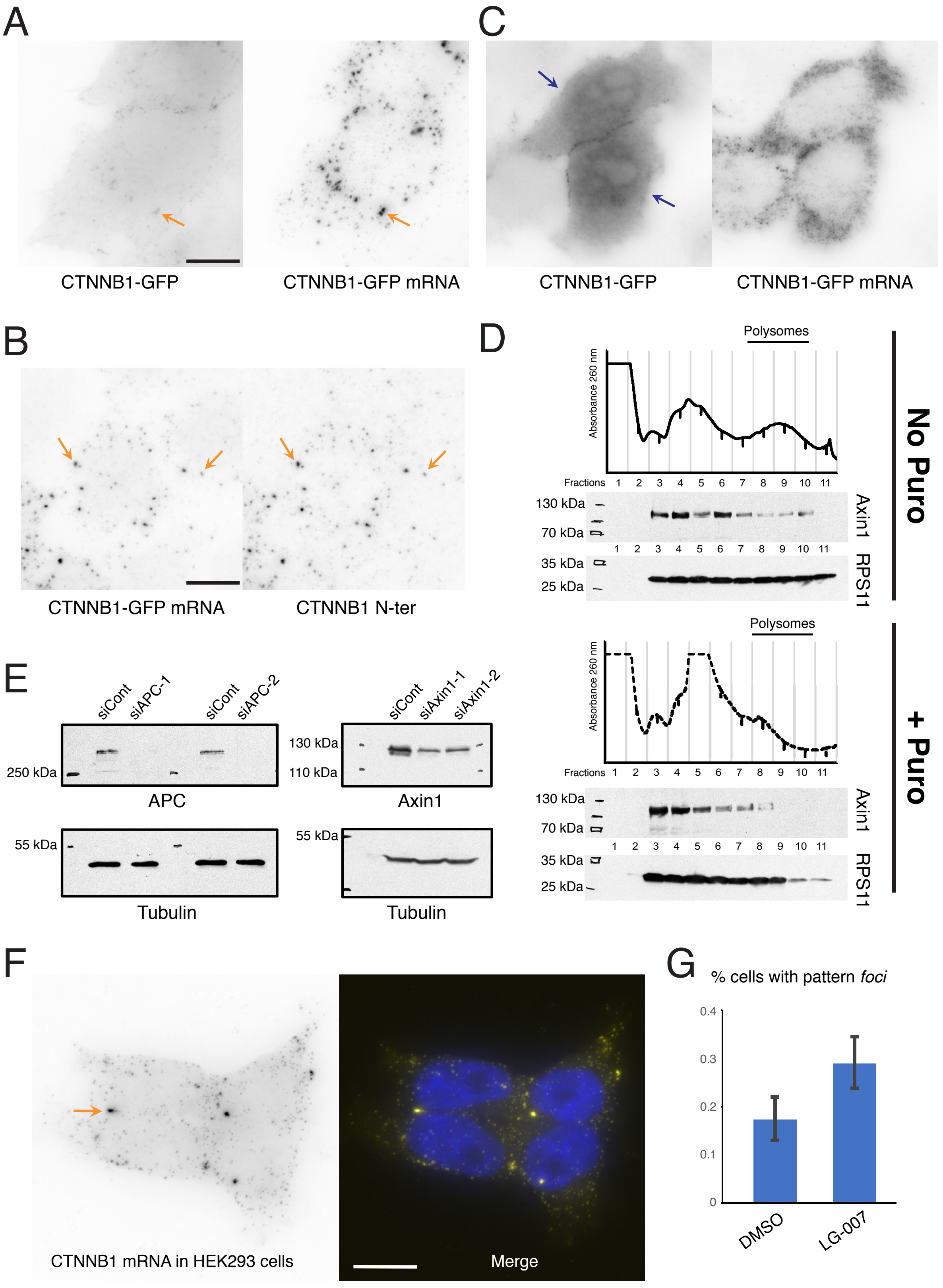
A
CTNNB1-GFP
CTNNB1-GFP mRNA
C
CTNNB1-GFP
CTNNB1-GFP mRNA
B
CTNNB1-GFP mRNA
CTNNB1 N-ter
D
Polysomes
Absorbance 260 nm
Fractions 1 2 3 4 5 6 7 8 9 10 11
130 kDa
70 kDa
35 kDa
25 kDa
Axin1
RPS11
No Puro
+ Puro
E
siCont
siAPC-1
siAPC-2
siAxin1-1
siAxin1-2
250 kDa
130 kDa
110 kDa
55 kDa
APC
Axin1
Tubulin
F
CTNNB1 mRNA in HEK293 cells
Merge
G
% cells with pattern foci
0
0.1
0.2
0.3
0.4
DMSO
LG-007

Figure S10

**Figure S10, related to Figure 9. β-catenin translation factories.**

A-C-Expression of β-catenin-GFP protein correlates with an absence of β-catenin mRNA foci. Legend as in Figure 1B, except the cells expressed a tagged CTNNB1 BAC. Orange arrow: β-catenin mRNA foci; blue arrow: accumulation of β-catenin-GFP protein in the nucleus. Scale bar: 10 microns.

B-β-catenin mRNA foci are labelled by an antibody against β-catenin N-terminus. Images are micrographs of HeLa cells expressing the tagged CTNNB1 BAC and both hybridized *in situ* with Cy3-labelled probes against the tag (panel CTNNB1-GFP mRNA), and incubated with Cy5-labelled anti-β-catenin antibody binding the N-terminus of the protein (left; panel CTNNB1-N-ter). Arrow point to foci. Scale bar: 10 microns.

D-Axin1 is present in polysomal fractions in a translation-dependent manner. The graphs depict the absorbance at 260nm in the different fraction of HeLa cells extract fractionated on a sucrose density gradient. The fraction containing polysomes is indicated. The images are Western blots analyses of the same fractions probed with the indicated antibodies. Molecular weight markers are indicated on the left. Top panels: untreated cells; bottom panels: cells treated for 1h with puromycin.

E-Efficacy of siRNAs against APC and Axin1. The images are Western blots of HeLa cells treated with the indicated siRNAs and probed with the indicated antibodies. Molecular weight markers are indicated on the left.

F-Endogenous β-catenin mRNAs accumulate in foci. Legend as in Figure S8A, except that probes bound endogenous β-catenin mRNA in HEK293 cells. Orange arrow points a mRNA foci. Scale bar: 10 microns.

G-The graph displays the percent of cells classified in the pattern *'foci'* by the automated classifier, after detecting endogenous β-catenin mRNA by smiFISH in HEK293 cells treated with LG-007 for 2 hours, or DMSO as control. Error bars: confidence interval computed by bootstrapping.
